# Structural variant evolution after telomere crisis

**DOI:** 10.1101/2020.09.29.318436

**Authors:** S.M Dewhurst, X Yao, Joel Rosiene, Huasong Tian, Julie Behr, Nazario Bosco, Kaori K. Takai, T de Lange, M Imielinski

**Affiliations:** Laboratory of Cell Biology and Genetics, Rockefeller University, New York, USA; Tri-Institutional Ph.D. Program in Computational Biology and Medicine, Weill Cornell Medicine, New York, USA; Department of Pathology and Laboratory Medicine, Englander Institute for Precision Medicine, and Meyer Cancer Center, Weill Cornell Medicine, New York, USA; New York Genome Center, New York, USA; Department of Biochemistry and Molecular Pharmacology, Institute for Systems Genetics, NYU Langone Health, New York, USA

## Abstract

Telomere crisis contributes to cancer genome evolution, yet only a subset of cancers display breakage-fusion-bridge (BFB) cycles and chromothripsis, hallmarks of previous experimental telomere crisis studies. We examine the spectrum of SVs instigated by natural telomere crisis. Spontaneous post-crisis clones from prior studies had both complex and simple SVs without BFB cycles or chromothripsis. In contrast, BFB cycles and chromothripsis occurred in clones that escaped from telomere crisis after CRISPR-controlled telomerase activation in MRC5 fibroblasts. This system revealed convergent evolutionary lineages altering one allele of 12p, where a short telomere likely predisposed to fusion. Remarkably, the 12p chromothripsis and BFB events were stabilized by independent fusions to 21. Telomere crisis can therefore generate a wide spectrum of SVs, and lack of BFB patterns and chromothripsis does not indicate absence of past crisis.

---

Structural variation is a hallmark of cancer genomes. Recent pan-cancer whole genome sequencing (WGS) studies have begun to reveal a more complete picture of the spectrum of structural variants (SVs) found in cancer genomes, ranging from simple deletions, duplications and translocations to complex and often multi-chromosomal rearrangements^1–3^. The PCAWG consortium catalogued WGS variants across >2,500 cases spanning 38 tumor types^4^ to nominate novel classes of complex SVs and cluster these into signatures, mirroring previous work in the categorization of single nucleotide variants (SNVs) into distinct mutational processes^5–8^. The analysis of genome graphs provides a rigorous and unified framework to classify simple and complex SVs (including chromothripsis, breakage-fusion-bridge (BFB) cycles, and double minutes), nominate novel event classes, and study the rearranged structure of aneuploid alleles^3^.

However, despite advances in the identification and classification structural variations, a mechanistic understanding of the underlying causes is often still lacking. SV mutational processes may have a more complex etiology than those driving the formation of SNVs and generate a more complex spectrum of patterns: layers of simple SVs can reshape a locus gradually and across multiple alleles, and complex SVs can rapidly rewire many genomic regions. In addition, multiple underlying causes can lead to the same type of rearrangement, and diverse outcomes can originate from a single cause. Further, it has been expensive and technically challenging to delineate specific mechanisms, although some progress has been made^9–14^.

Telomere crisis, which is thought to occur at an early stage of carcinogenesis before a telomere maintenance mechanism is activated^15^, has been nominated as a cause of cancer genome SVs. *A priori*, the genomic consequences of telomere crisis are predicted to be profound: critically short telomeres in human cells can trigger a DNA damage response, and inappropriately engage DNA repair pathways resulting in telomere to telomere fusions^16,17^. Subsequent cell divisions in the presence of fused chromosomes has long been considered a mechanism driving complex chromosomal rearrangements such as BFB cycles in tumors^18,19^. The characteristic fold-back inversions of BFB cycles are known to contribute to tumorigenesis in ALL^20^ as well as squamous cell cancers and esophageal adenocarcinoma^3^. Modeling of telomere crisis in late generation telomerase-deficient mice lacking p53 showed that telomere dysfunction engenders cancers with non-reciprocal translocations, as well as focal amplifications and deletions in regions relevant to human cancers^21,22^. Furthermore, mouse models of telomerase reactivation after a period of telomere dysfunction showed that acquisition of specific copy number aberrations and aneuploidy could drive malignant phenotypes^23^.

Studies in cultured human cells have also illuminated the genomic consequences of telomere dysfunction. Even a single artificially deprotected telomere can fuse with multiple intra- and inter-chromosomal loci leading to complex fusion products^24^ but it is unclear whether these complex rearrangements are compatible with escape from telomere crisis. The resolution of dicentric chromosomes induced by over-expression of a dominant negative allele of the telomere binding protein TRF2 can lead to the dramatic chromosome re-shuffling phenomenon of chromothripsis^11,14,25^. However, to date the only study directly investigating the consequences of a sustained period of telomere dysfunction failed to identify any complex rearrangements in wild-type HCT116 colon carcinoma cells^26^. This may be because these cells readily escaped from the telomere dysfunction induced by expression of a dominant-negative hTERT (telomerase reverse transcriptase) allele. In HCT116 cells with defective Non-Homologous End Joining (NHEJ) pathways, which are already genetically unstable, complex chained SVs were observed after telomere dysfunction, but the relevance of these types of rearrangements to human cancer remains unclear^26^.

Given the expanding repertoire of structural variation present in so many cancer types, and the potential contribution of telomere dysfunction to many of these different aberrations, we set out to characterize the extent and type of structural variation that can be unleashed by telomere crisis and subsequent genome stabilization by telomerase expression. We approached this problem in two ways. First, we performed whole genome sequencing (WGS) on a panel of seven previously isolated cell lines that had escaped telomere crisis spontaneously through telomerase activation. Secondly, we created a controlled *in vitro* telomere crisis system by engineering an MRC5-derived cell line in which telomerase could be activated during telomere crisis and analyzed the resulting post-crisis clones by WGS. The results indicate that the consequences of telomere crisis are varied and not readily predictable. In many instances, the post-crisis genomes did not feature BFB cycles or chromothripsis. Therefore, telomere crisis may have occurred in cancers that lack these genomic scars of telomere dysfunction.

## Results

### Genomic complexity after spontaneous telomerase activation

In order to determine the SVs in post-telomere crisis genomes, we examined seven SV40 large T-transformed cell lines that had undergone spontaneous telomerase activation after passage into telomere crisis (Supplementary Table 1, Supplementary Figure 1). The cell lines represent independent immortalization events in a variety of cell lineages^27–29^. We carried out whole genome sequencing of these cell lines to a median depth of 41X (range: 17-55) and generated junction balanced genome graphs^3^ via JaBbA from SvABA^30^ and GRIDSS^31^ junction calls (see Methods). All genomes demonstrated some level of aneuploidy but there was a high variance in the total number of junctions fit across each genome, ranging from 7 to 100. Analysis of junction-balanced genome graphs^3^ revealed complex multi-chromosomal gains in four of seven samples, with the remaining three lines harboring only broad arm level losses (Figure 1A).

**Figure 1.**
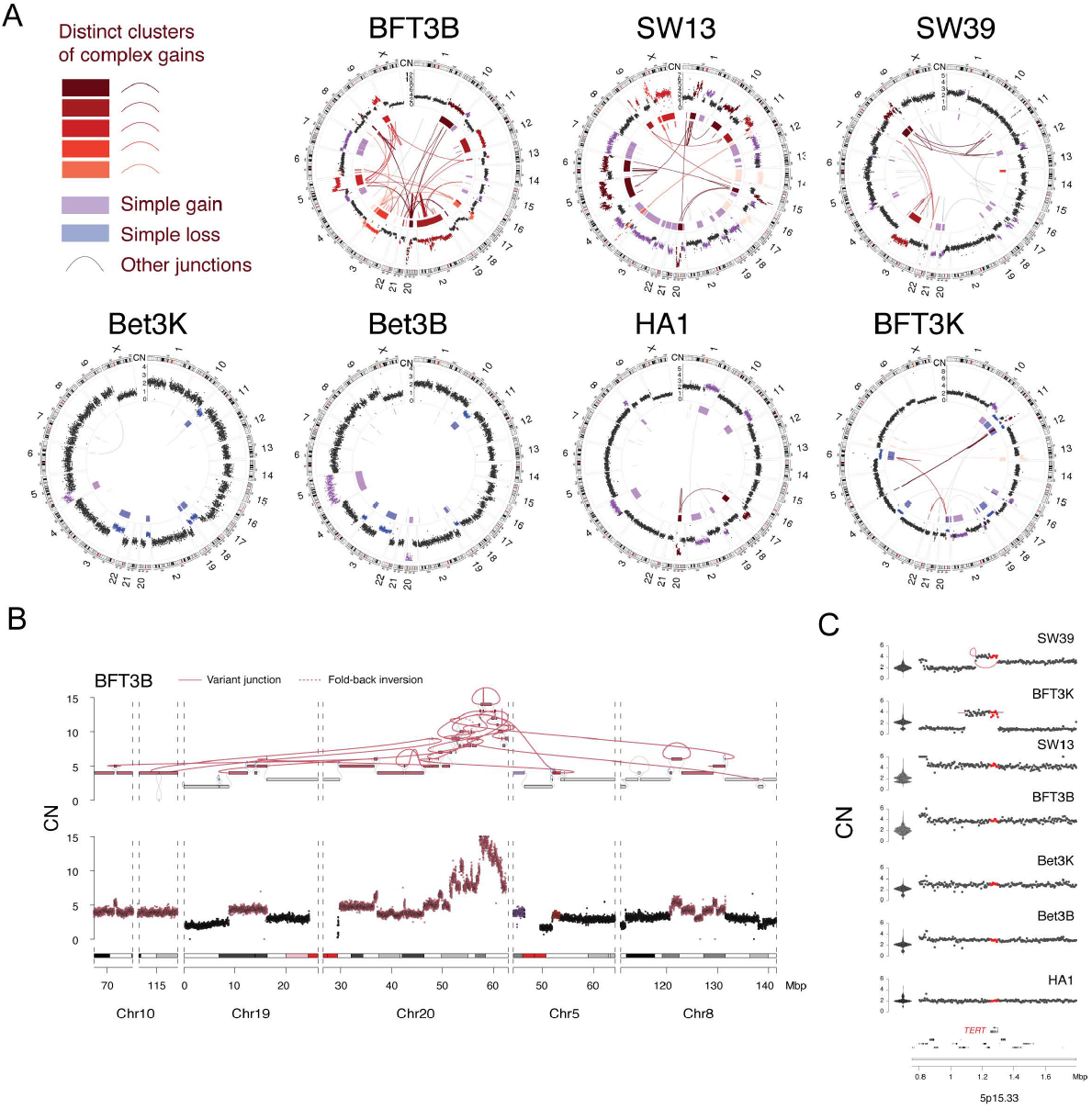
Genomic complexity after spontaneous telomerase activation. **A)** CIRCOS plots showing seven cell lines that emerged spontaneously from telomere crisis (Supplementary Table 1), four of which show one or more clusters of complex gains. Binned purity- and ploidy-transformed read depth is shown in the periphery, with colored links in the center representing variant (rearrangement) junctions. A series of red colors is used to show junctions and read depth bins belonging to distinct clusters of complex gains in each cell line. Additional colors describe junctions and bins, including those belonging to simple losses and gains. See Methods for additional details regarding junction and bin classifications. **B)** An example of a complex gain cluster in cell line BFT3B, spanning six discontiguous regions across five chromosomes, including a focus of high-level amplification amplifying telomeric portions of chr. 20p to 15 copies. Bottom track showing binned purity- and ploidy-transformed read-depth, with the top track showing the associated junction-balanced genome graph (see Methods^3^) with y-axis representing units of per cell copy number (CN) across bins and graph nodes (i.e. intervals). Gray and colored edges represent reference and variant junctions, respectively. Blue edges represent loose ends (see Methods for further details). Bins and junctions are colored as in panel A. **C)** Read depth and junction patterns at the *TERT* locus across the seven post-crisis clones. Each track shows binned purity- and ploidy-transformed read depth in units of CN, with variant (rearrangement) junctions and loose ends plotted as red arcs. The bottom track highlights the *TERT* gene among genes on chromosome 5p15. Violin plots to the left of tracks show the genome-wide distribution of read depth in units of CN, demonstrating that 6 of 7 clones have elevated CN at the *TERT* locus.

Strikingly, genome graph-based categorization of complex SVs^3^ did not identify the classic footprints of chromothripsis or BFB cycles in these genomes. However, several amplified subgraphs were associated with stepwise copy number gains reminiscent of BFB cycles (Figure 1A-B, Supplementary Figure 1). The majority of copy changes in these subgraphs could not be attributed to fold-back inversion junctions (a hallmark of BFB cycles) but were instead driven by a spectrum of duplication and translocation-like junctions and templated insertion chains. These patterns are exemplified in a 10 Mbp region of 20q of BFT3B that is amplified to 10-15 copies, incorporating Mbp scale fragments of chromosomes 5p, 8q, 10q, as well as 19p at lower copy number (Figure 1B). Of note, six of seven cell lines showed modest increases in *TERT* copy number, providing a possible genomic basis for telomere crisis escape (Figure 1C). Among these, the four cell lines harboring the most complex genome graphs had the highest *TERT* copy number, although *TERT* was not among the most complex or highly amplified loci in these lines. Across seven different transformed cell lines, spontaneous escape from telomere crisis was associated with a highly variable spectrum of SV patterns comprising complex and non-canonical patterns of amplification and numerical gains and losses, including some affecting *TERT*.

### An *in vitro* system for telomerase-mediated escape from natural telomere crisis

To gain a clearer insight into the nature of SVs that arise during telomere crisis, we developed an *in vitro* system in which we could reproduce telomere crisis and generate a high number of post-crisis clones. MRC5 human lung fibroblasts were chosen to model telomere crisis since they have a well-defined *in vitro* replicative potential determined by telomere attrition. To bypass senescence, the Rb and p21 pathways were inactivated by infecting the population of MRC5 cells with a retrovirus bearing shRNAs targeting the respective transcripts (Supplementary Figure 2A). This population of MRC5/Rbsh/p21sh was then endowed with an inducible CRISPR activation system (iCRISPRa) to activate the *TERT* promoter (Supplementary Figure 2B). The iCRISPRa system employed a doxycycline-inducible nuclease-dead Cas9 fused to a tripartite transcriptional activator (VP64-p65-Rta)^32^ and four gRNAs targeting the *TERT* promoter (Figure 2A, Supplementary Figure 2B). Addition of doxycycline (dox) to MRC5/Rbsh/p21sh/iCRISPRa-TERT cells resulted in robust induction of *TERT* mRNA within 96 hours, whereas without dox, *TERT* transcripts are undetectable in this cell line (p<0.001, Figure 2B). A similar dox-induced increase in mRNA expression was noted upon introduction of sgRNAs to a control gene (Supplementary Figure 2C). Induction of telomerase activity was readily detectable in a TRAP (telomerase repeated amplification protocol) assay (Figure 2C). However, the induced *TERT* mRNA levels and the TRAP activity were significantly lower than in telomerase-positive control cell lines. The relatively weak telomerase activity in this system harmonizes with recent work showing that *TERT* promoter mutations initially result in low levels of telomerase activity that is not sufficient to maintain bulk telomere length^33^.

**Figure 2.**
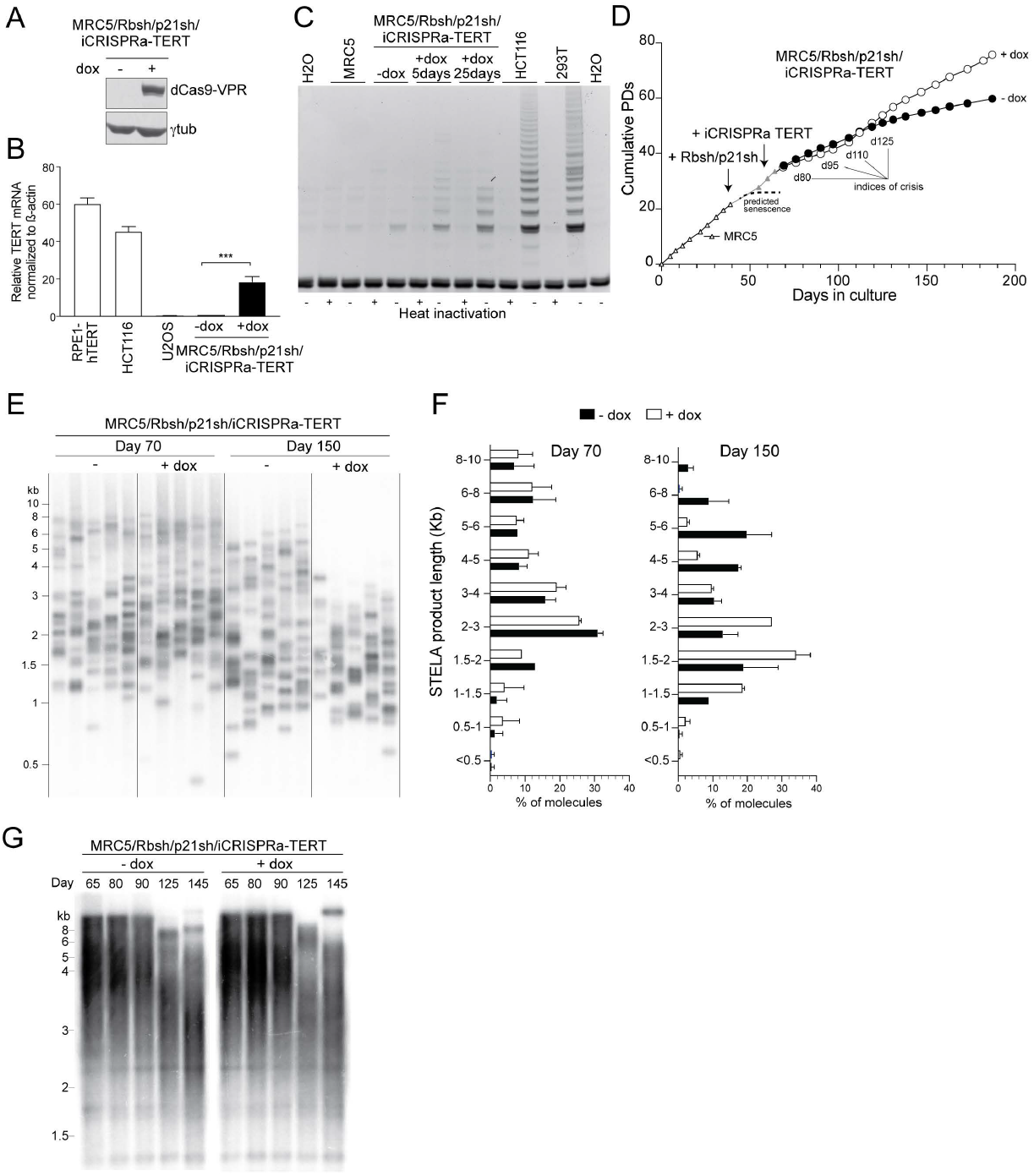
An *in vitro* system for telomerase-mediated escape from natural telomere crisis. **A)** Immunoblot for dCas9-VPR (using a Cas9 Ab) in MRC5/Rbsh/p21sh/iCRISPRa-TERT cells with or without doxycycline treatment for 96 h (see also Supplementary Figure 2A-B). **B)**qPCR of TERT mRNA expression in RPE1, HCT116, U2OS and MRC5/Rbsh/p21sh/iCRISPRa-TERT cells with and without doxycycline treatment. Values are normalized to *β*-actin mRNA. Data from 3 independent biological replicates; error bars show means ±SDs. **C)** TRAP assay on extracts from MRC5 and MRC5/Rbsh/p21sh/iCRISPRa-TERT cells with and without doxycycline treatment for indicated time periods. HCT116 and 293T (Phoenix) cells are included as positive controls. **D)** Growth curve of parental MRC5 cells, MRC5/Rbsh/p21sh cells, and MRC5/Rbsh/p21sh/iCRISPRa-TERT cells grown with and without doxycycline. Arrows indicate when each construct was introduced. Days in culture represents total time in culture from parental MRC5 cells to late passage MRC5/Rbsh/p21sh/iCRISPRa-TERT cells. Time points for telomere analysis (presented in Figure 3) and the approximate onset of senescence in the parental MRC5 cells are indicated. **E)** STELA of XpYp telomeres in MRC5/Rbsh/p21sh/iCRISPRa-TERT cells with or without doxycycline treatment at 70 and 150 days of culture. **F)** Quantification of band intensity in E, with background signal subtracted. Data from two independent biological experiments is shown (see also Supplementary Figure 2D). Biological replicates represent cells at approximately the same days in culture (± 5 days). **G)** Genomic blot of telomeric *Mb*oI/*Alu*I fragments in MRC5/Rbsh/p21sh/iCRISPRa-TERT cells grown with or without doxycycline at the indicated time points.

At approximately 120 days after the start of the experiment (55 days with dox) the MRC5/Rbsh/p21sh/iCRISPRa-TERT population was proliferating faster than their untreated counterparts (Figure 2D). Inspection of individual telomere lengths using Single Telomere Length Analysis (STELA^34^) revealed that although telomerase expression was sufficient to allow the cells to proliferate, it was not sufficient to maintain bulk telomere length (Figure 2E). After 150 days of continuous culture, the majority (86%) of XpYp telomeres in induced MRC5/Rbsh/p21sh/iCRISPRa-TERT cells were between 1-4 kb compared to 40% in uninduced cells (Figure 2F, Supplementary Figure 2D). Consistent with this, genomic blotting showed bulk telomere shortening in both induced and uninduced cells (Figure 2G). These telomere dynamics are consistent with the expectation that in the culture without telomerase, cells with critically short telomeres will preferentially be lost, leading to a surviving population with relatively longer telomeres. In contrast, cells in the induced culture with (low) telomerase activity have the ability to elongate the shortest telomeres. As a result, the induced cells will tolerate telomere attrition better and present with overall shorter telomeres at later time points.

### Dissipating telomere crisis in MRC5/Rbsh/p21sh/iCRISPRa-TERT cells

To confirm that the MRC5/Rbsh/p21sh/iCRISPRa-TERT cells experienced telomere crisis before the induction of telomerase increased their proliferation rate, we investigated cells at various time points from the start of the experiment (Figure 2D). Metaphase spreads showed both induced and uninduced MRC5/Rbsh/p21sh/iCRISPRa-TERT cells contained dicentric and multicentric chromosomes (Figure 3A) and genomic blots showed high molecular weight telomere bands consistent with fused telomeres (Figure 2G). As expected from the ability of telomerase to counteract the formation of critically short telomeres, at 125 days after the start of the experiment, induced cells had significantly fewer fusions than untreated cells (21% vs. 40%, p<0.05, Figure 3B, Supplementary Figure 3A). PCR-mediated detection of fusions between the Tel Bam 11 family of telomeres^35,36^ confirmed these dynamics (Figure 3C). Quantification of the fusion frequency showed a significant reduction in number of fusions per haploid genome in the induced population (day 110, p<0.01, Figure 3D). Consistent with telomerase-mediated genome stabilization, there was a trend towards a lower level of 53BP1-marked DNA damage foci at later time points (Figure 3E-F) and the percentage of cells with micronuclei (an indicator of genome instability) was significantly reduced at day 110 (p<0.05, Figure 3G). Taken together, these data indicate that after a period of genomic instability induced by critically short telomeres, iCRISPRa-mediated telomerase activation is sufficient to stabilize the genome and allow the MRC5/Rbsh/p21sh/iCRISPRa-TERT cells to escape from telomere crisis.

**Figure 3.**
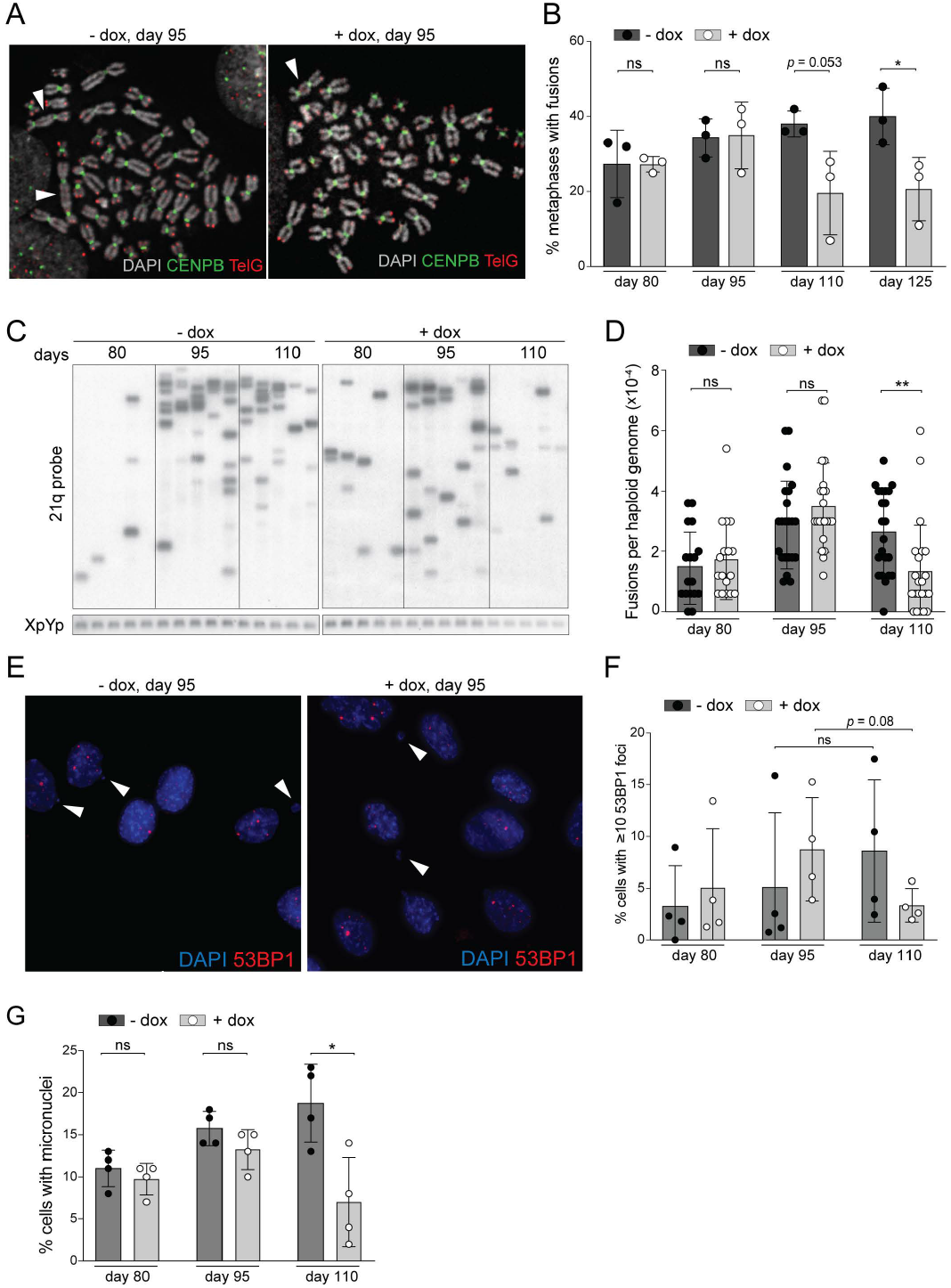
Dissipating telomere crisis in MRC5/Rbsh/p21sh/iCRISPRa-TERT cells. **A)** Metaphase spreads from MRC5/Rbsh/p21sh/iCRISPRa-TERT cells with and without doxycycline at day 95. Telomeres are detected with a telomeric repeat PNA probe (TelG, red) and centromeres are detected with a probe for CENPB (green). DNA was stained with DAPI (gray). Chromosome fusions are indicated by white arrowheads. **B)** Quantification of percentage of metaphase spreads with at least one fusion after the indicated days of continuous culture for MRC5/Rbsh/p21sh/iCRISPRa-TERT cells with and without doxycycline. Error bars indicate means and SDs from three independent biological replicates. *P* values were determined with an unpaired student’s t-test; ns, significant, *, *p* <0.05 (see also Supplementary Figure 3A). **C)** Gel showing products of telomere fusion PCR on MRC5/Rbsh/p21sh/iCRISPRa-TERT cells cultured with and without doxycycline for the indicated time. Each lane represents an independent replicate PCR reaction. Telomere fusion products are detected by hybridization with a probe for the 21q telomere (see Methods) and the control XpYp PCR product is detected with Ethidium bromide staining. **D)** Quantification of the number of telomere fusion products per haploid genome using the assay shown in panel C. Each dot represents a single PCR reaction. Reactions from two independent biological replicates are shown. *P* values were determined as in panel B. **, *p* <0.01. **E)** Detection of micronuclei (arrowheads) and DNA damage foci using indirect immunofluorescence for 53BP1 (red) in the indicated cells. DNA is stained with DAPI (blue). **F)** Quantification of percentage of cells with *≥*10 53BP1 foci at the indicated time points. Data show the means from 3 independent biological replicates with SDs. *P* values as above. **G)** Quantification of percentage of cells with micronuclei after indicated days in culture. Data show the means from 3 independent biological replicates with SDs. P values as above.

### Genomic screening of post-crisis clones

To assess the genome structure of proliferating post-crisis cells, single cell clones were isolated from induced MRC5/Rbsh/p21sh/iCRISPRa-TERT cells at day 120 (‘Y clones’) and day 150 (‘Z clones’) (Figure 4A). The clonal yield at day 150 was greater than at day 120 in induced cells but no clones could be isolated from the uninduced population at either timepoint. The lower clonal yield at day 120 may be due to incomplete stabilization of the telomeres since clones from this time-point showed a higher burden of fused telomeres than those derived from day 150 (Supplementary Figure 4A). Post-crisis clones from both timepoints showed evidence of ultra-short telomeres and reduced telomere length (Supplementary Figure 4B-C). Telomerase activity in post-crisis clones was comparable to the parental induced population, indicating that clone viability was not due to selection for increased telomerase activity (Supplementary Figure Figure 4D). To generate control clones which had not passed through a period of telomere crisis, early passage MRC5 cells were infected with a retrovirus expressing hTERT and single cell clones were isolated (Supplementary Figure 4E). Genome profiling with low pass (∼5X) WGS was performed on eight hTERT-expressing control clones (CT clones), 36 Y clones from day 120, and 82 Z clones from day 150 (Table S2).

**Figure 4.**
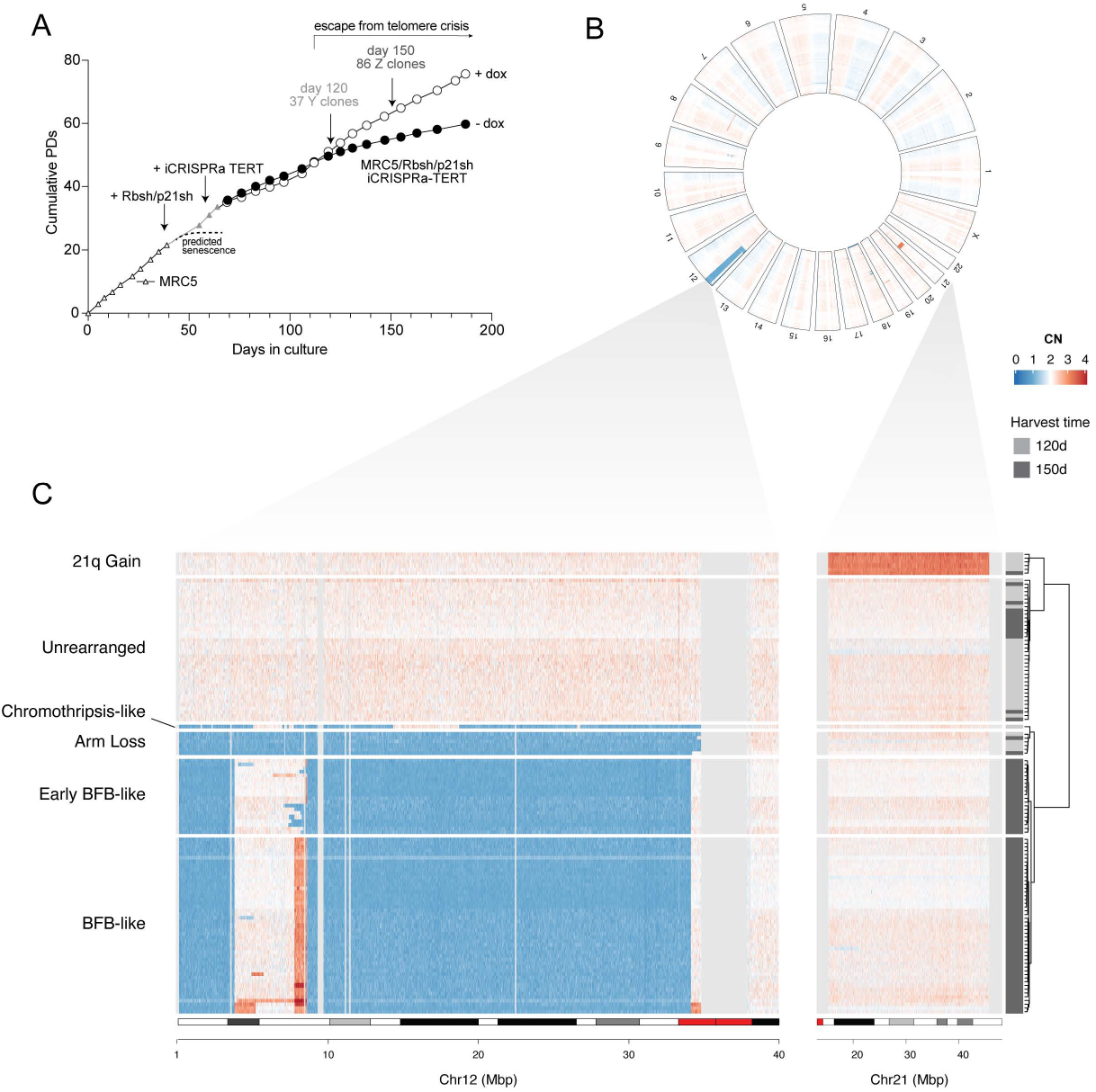
Genomic screening of post-crisis clones. **A)** Growth curve of MRC5/Rbsh/p21sh/iCRISPRa-TERT cells with and without doxycycline, indicating the time points at which single cell clones were derived (day 120 and day 150). **B)** Circular heatmap showing genome-wide binned purity- and ploidy-transformed read depth (in units of CN across 118 low pass WGS-profiled clones. Heatmap rows correspond to concentric rings in the heatmap. Clones are clustered with respect to genome-wide copy number profile similarity (see Methods). **C)** Zoomed in portion of chromosomes 12 and 21 that underwent copy number alterations in a majority of the clones, clustered based on their coverage across these regions. Clusters are named with respect to their consensus copy number pattern, and on the basis of high depth WGS analyses presented in Figure 5. Chromosome 21 gain n=6 clones, unrearranged n=38, chromothripsis-like n=1, arm loss n=6, early BFB-like n=20, BFB-like n=47.

Analysis of genome-wide read depth across 118 Y and Z clones demonstrated predominantly diploid genomes with a striking enrichment of clones with DNA loss on most of chromosome 12p (63%, 74/118, Figure 4B). Within the other 44 samples, we observed a subset of clones (5%, 6/118) with gains of chromosome 21q. As expected, control CT clones showed no evidence of SVs or copy number variants (Supplementary Figure 4F). Hierarchical clustering of all clones by their coverage on chromosomes 12p and 21q revealed six distinct clusters (Figure 4C). A minority of clones were diploid on chromosomes 12 and 21 and elsewhere in the genome and are therefore designated as ‘unrearranged’ (32% of clones, 38/118). Of note, the unrearranged group was enriched in day 120 samples (*P* = 1.79⨯10^−9^, odds ratio 14.7, Fisher’s exact test, Figure 4C), suggesting that these clones may not have experienced crisis prior to telomerase induction. The cluster of clones with 21q gain were diploid on 12p.

The remaining 74 clones (63%) all showed a heterogenous pattern of copy number alterations targeting 12p (Figure 4C). First, a singular pattern of distinct interspersed losses that resembled chromothripsis was present in one clone (0.8%, 1/118). Another cluster of 6 clones demonstrated complete loss of 12p (‘arm loss’, 5%, 6/118). The other 67 clones all share a breakpoint near the distal end of 12p and a large deletion starting ∼9 Mbp from the centromere. These clones were differentiated into two clusters by the presence or absence of an amplification around 8∼9 Mbp from the 12p telomere. In the 47 clones that contained this amplification, by aggregating a consensus read depth profile we observed stepwise gains at the distal end of 12p, a pattern reminiscent of BFB cycles (Supplementary Figure 5A). This cluster was therefore labelled ‘BFB-like’, a designation which is further supported by data presented below. The 20 clones (17%) that lack the amplicon around 8-9 Mbp harbored varying boundaries of the shared larger deletion, and based on the analysis described below we designate these as ‘early BFB-like’. In summary, these low-pass WGS copy number profiles indicated a limited set of distinct lineages surviving telomere crisis, with at least two lineages independently converging on chromosome 12.

**Figure 5.**
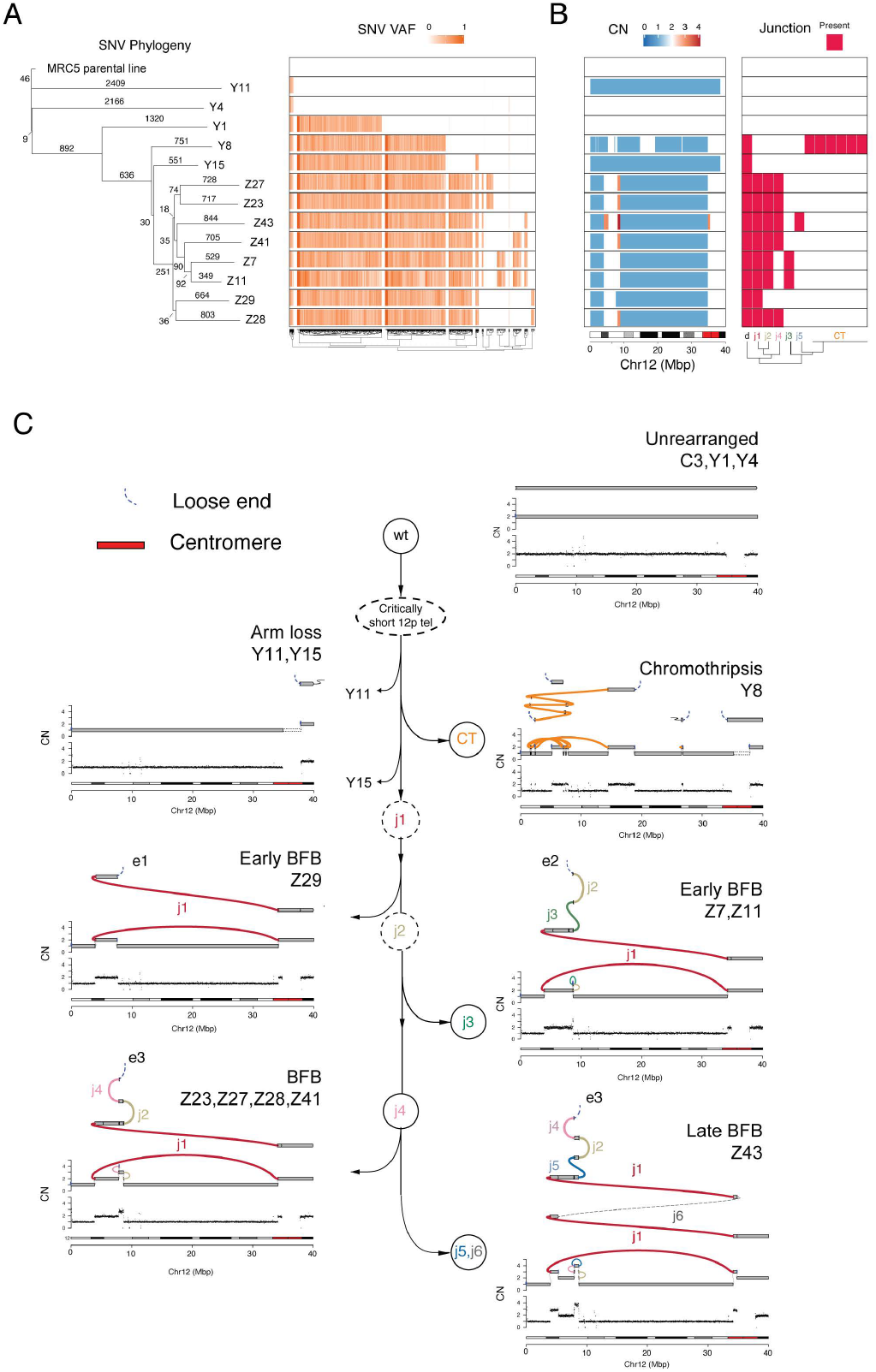
High-resolution reconstruction and lineage of post-crisis genomes. **A)** SNV-based phylogeny inferred across 13 high-depth WGS clones and heatmap of variant allele fractions (VAF) for SNVs detected among two or more clones. For simplicity, private SNVs (those found only in a single clone) are not shown. **B)** Heatmap of chromosome 12p copy numbers and variant junction patterns in chromosome 12 (see text and Methods). **C)** Tree showing distinct trajectories of structural variant evolution following 12p attrition and subsequent telomere crisis. Each terminal node in in the tree is associated with a unique 12p profile comprising a representative binned read depth pattern (bottom track) from one or more clones mapping to an identical junction-balanced genome graph (second track from bottom). The top track in each profile represents a reconstruction of the rearranged allele. Each allele is a walk of genomic intervals and reference/variant junctions that, in combination with an unrearranged 12p allele (not shown), sums to the observed genome graph (see Methods). Two distinct arrows linking Y11 and Y15 demonstrate that these clones are distinct lineages (based on divergent SNV patterns, see panel A), that converge to identical WGS 12p CN profiles (although with likely distinct breakpoints inside the 12p centromere unmappable by WGS, see text).

### High-resolution reconstruction and lineage of post-crisis genomes

To gain further insight into structural variant evolution along these lineages, we chose 15 representative clones spanning the 5 clusters with rearrangements involving 12p for high-depth WGS to a median read depth of 50X (range: 30-88). Phylogenies derived from genome-wide SNV patterns demonstrated a median branch length of 551 SNVs (range: 9-2,409), a low mutation density (<1 SNV/Mbp) that is consistent with previous WGS studies of clones in cell culture^37^. This analysis revealed four major clades (Figure 5A). These clades had good concordance with copy number alteration and rearrangement junction patterns in the same 12p region, suggesting these clones represent distinct post-crisis evolutionary lineages (Figure 5B).

In order to further reconcile the shared and distinct rearrangement junctions present in the evolution of these clones, we carried out local assembly of rearrangement junctions and junction balance analysis (see Methods^3^), which revealed 7 distinct junction-balanced genome graphs spanning 12p (Figure 5C). With the exception of the chromothriptic lineage (see below), each of these distinct lineages was represented by more than one post-crisis clone. We then applied gGnome to infer a set of linear alleles parsimoniously explaining the different genome graph patterns^3^ (see Methods, Figure 5C).

Analyzing the clonal evolution of these rearranged 12p alleles, we identified 8 clones demonstrating progressive stages of a BFB cycle. This complex variant evolved after a long-range inversion junction (j1) joined a distal end of 12p to its peri-centromere. This junction was followed by subsequent fold back inversion junctions (j2, j3, j4), clustered at the 8-9 Mbp focus on 12p, which are present in two different sets of post-crisis clones (Early BFB, BFB, Figure 5C). The earliest of the fold-back inversion junctions (j2) in the BFB lineage was associated with a cluster of 3 G or C mutations within 2 kbp of each other, consistent with APOBEC-mediated mutagenesis^25^ (Supplementary Figure 5B). The most complex locus in the BFB lineage (Z43, Late BFB, Figure 5C), contained six variant junctions in *cis*, including two late tandem duplications (j5, j6). Although j6, which connects the distal portion 12p to the 12p centromere, was not directly observed in the short read WGS data, it was imputed (dashed line, j6, Figure 5C) to resolve the duplication of j1 in clone Z43, as well as two allelic ends in the genome graph. Remarkably, the vast majority (97%) of SNVs detected in this BFB lineage (Figure 5A) were either shared or private, indicating that these stages of BFB evolution occurred rapidly in the history of the experiment.

We confirmed a chromothripsis event in an independent lineage (Y8), which lacked j1 and all subsequent junctions of the BFB lineage, further supporting the idea that this is an independent lineage (Figure 5C). Integration of SCNA data with the SNV phylogeny showed clones from the unrearranged lineage (Y1 and Y4) and one of the 12p arm loss clones (Y11) to be mutationally distant (>2,000 SNVs) from the chromothripsis (Y8) and BFB lineages, which shared over 1,583 SNVs (Figure 5A). Supporting this, a small (∼21.5 Kbp) simple deletion junction was shared across Y8, Y15, and all the BFB lineage samples, yet was absent in Y11 (Supplementary Figure 5C).

This comparison established that the 12p loss in Y11 could not have occurred after j1, and indicates that a second independent arm loss must have given rise to Y15. Interestingly, the Y15 arm loss clone was clustered in the BFB/Y8 clade in the SNV phylogeny, sharing 30 SNVs with the BFB lineage which it did not share with Y8 (Figure 5A). This indicates that the 12p arm loss in Y15 may have arisen either before or after j1. Although the breakpoints of the Y11 and Y15 arm losses could not be mapped due to their location in the 12 centromeric region, based on the SNV phylogeny, they likely represent distinct events. Taken together, these results support a model whereby at least three lineages independently rearranged a previously wild type 12p during telomere crisis (Figure 5C). Our data appear to have captured sequential steps in the formation of an increasingly complex BFB-like event. Each of these stages must represent a stabilized allele since the post-crisis lines are clonal, and multiple clones share the same rearrangement junctions (Figure 5B). This necessarily raises the question as to what caused the on-going instability, and how and where these complex alleles are terminated.

### Resolution of BFB cycles in telomere crisis

Analysis of junction-balanced genome graphs allows for the nomination of ‘loose ends’ (or allelic ends), representing copy number changes that cannot be resolved through assembly or mapping of short reads. We identified three distinct loose ends across the 4 variant graphs spanning the 8 clones in the BFB lineage (Figure 5C, Supplementary Figure 5D). Each of these loose ends were placed at the terminus of their respective reconstructed allele, and we posit they represent the new ‘ends’ of the derivative alleles of the BFB lineage. Distinct ends for each of these rearranged lineages suggests the derivative 12p allele could have been stabilized independently. We did not observe telomere repeat-containing reads mated to these loose ends, arguing against telomere healing at these loci. Instead, loose reads represented highly repetitive unmappable sequences which may be a result of the junctions being in close proximity to centromeric regions (see below).

To resolve the genomic architecture at these loci we generated karyotypes from metaphase spreads for representative rearranged clones (Figure 6A, Supplementary Figure 6A), which revealed that in the BFB and chromothripsis (Y8) lineages, the chromosome 12 derivative was linked to a copy of chromosome 21 with an intact long arm (Supplementary Figure 6A). These results were confirmed with chromosome painting, demonstrating a derivative chromosome transitioning between 12 and 21 (Figure 6B-C, Supplementary Figure 6B). Two possible events can explain these findings: the 12-21 fusion could have occurred as an early event during telomere crisis, preceding the divergence of Y8 (chromothriptic) and the BFB lineage; alternatively, independent 21 fusion events stabilized the derivative chromosome 12 following formation of the distinct junction lineages in Figure 5C. We consider the first possibility unlikely since the creation of the long-range inversion (j1) and subsequent fold back junctions in the BFB lineage would require the formation of interstitial 12p breaks on a 12p-21 derivative chromosome. Such breaks are predicted to result in the loss of 21, which would be distal to these junctions on the fusion allele. Furthermore, the acrocentric nature of chromosome 21 would make it more likely to stabilize the overall chromosome architecture, suggesting that an early 12-21 derivative chromosome would be unlikely to engage in the additional SV events observed in the BFB lineage. We therefore consider it likely that each of the BFB cycles and chromothripsis clones were independently resolved through subsequent fusion to 21.

**Figure 6.**
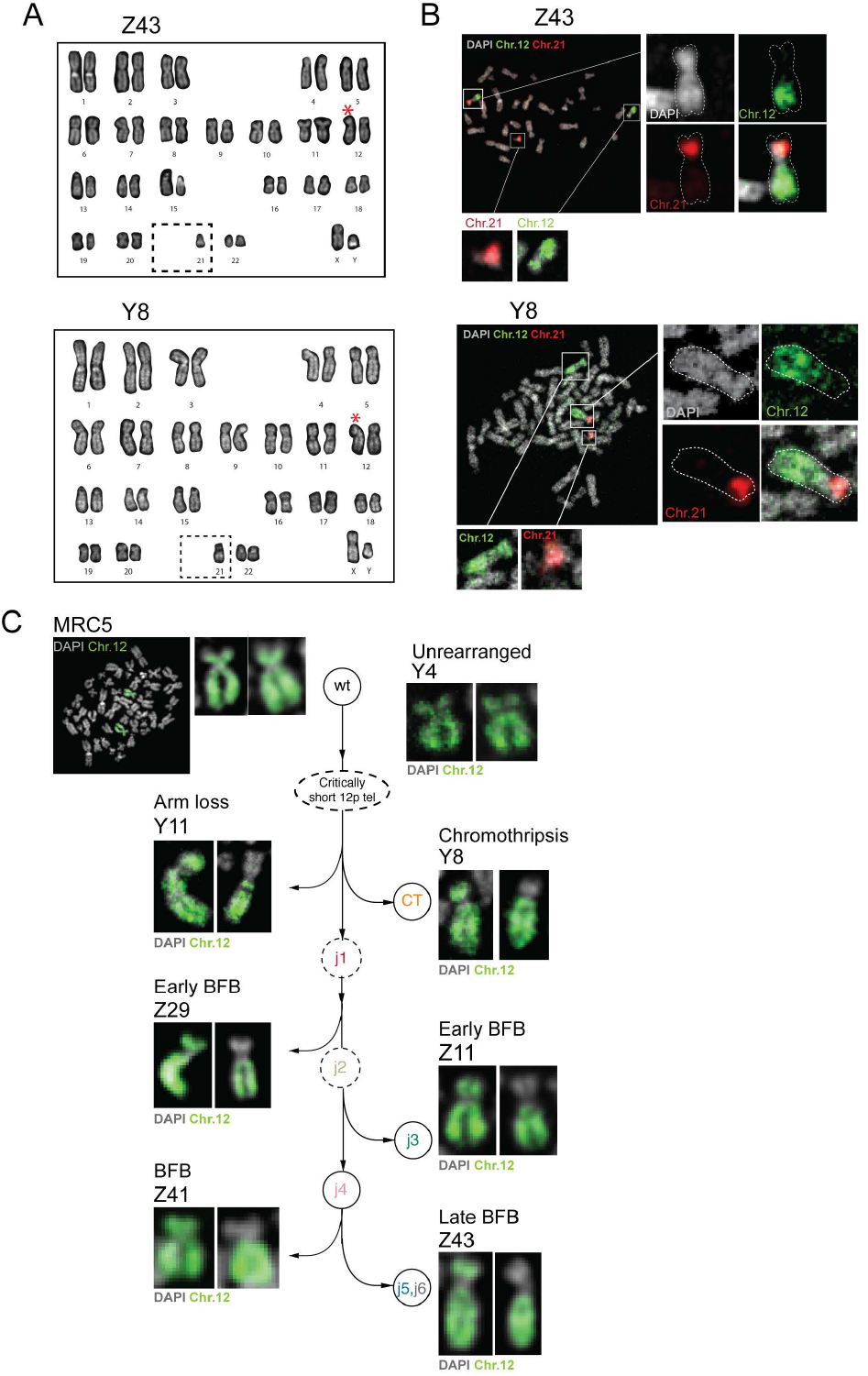
Resolution of BFB cycles in telomere crisis. **A)** DAPI banded karyotype of post-crisis clone Z43 and Y8 showing a rearranged chromosome 12 and loss of one copy of chromosome 21. Red star: Marker chromosome 12. Dashed gray box: absence of 21. (See also Supplementary Figure 6A). **B)** Representative metaphase spreads of clone Z43 and Y8 hybridized with whole chromosome paints for chromosomes 12 (green) and 21 (red). DNA was stained with DAPI (gray). Insets show enlarged images of the 12-21 derivative marker chromosome, and intact copies of the sister alleles (see also Supplementary Figure 6B). **C)** Images of chromosome 12 from representative clones from each branch of the evolution of chromosome 12 post-crisis (according to the analysis in Figure 5C). Metaphases were hybridized with whole chromosome 12 paint. DNA was stained with DAPI (gray).

Unlike the BFB and chromothripsis clusters, the two 12p arm loss lineages (Y11 and Y15) did not fuse to chromosome 21. In the Y11 clone, the derivative chromosome 12 appears to contain a distinct fusion (with a longer p-arm), consistent with it being an independent lineage (Figure 6C). We were unable to further resolve the nature of the stabilization events in these two clones. It would be necessary to perform long molecule DNA sequencing across different lineages in order to confirm the distinct nature of the fusion junction in each of the post-crisis clones.

### Telomere attrition renders 12p vulnerable

The convergent evolution patterns observed in our system suggests either 12p vulnerability to rearrangements or selection for 12p loss during telomere crisis. We believe strong selection is unlikely, given the existence of day 150 clones with diploid 12p (15.8%, 13/82, with or without 21q gain). The preferential rearrangement of the short arm of chromosome 12 in the post-crisis system could be explained if one of the two 12p telomeres is among the shortest telomeres in the MCR5 parental cells. Attrition of the shortest telomeres is predicted to generate the first telomere fusions and associated rearrangements in the culture.

We first asked whether the same parental allele was targeted across the chromosome 12-associated events in our cohort. Such allele specificity would argue against a selection for loss of 12p sequences as the driver for the 12p events since such selection should have occurred without preference for one allele. We phased heterozygous SNPs on 12p on the basis of whether they belonged to the lost (L) or retained (R) allele on the early 12p arm loss clone, Y11 (Figure 7A). Analyzing phased SNP patterns across all the high– and low-pass MRC5 clone WGS profiles in our dataset demonstrated that the L allele of 12p was the exclusive target of all chromosome 12 structural variants (Figure 7B, Supplementary Figure 7A). This included the clones from the chromothripsis (Y8), and BFB (Z43) lineages (Figure 7C), which our phylogenetic analyses associated with independent alterations on a previously unrearranged chromosome 12. On the basis of these results, we concluded that the short arm of the L allele of 12 was the most vulnerable to rearrangement in the MRC5 parental line.

**Figure 7.**
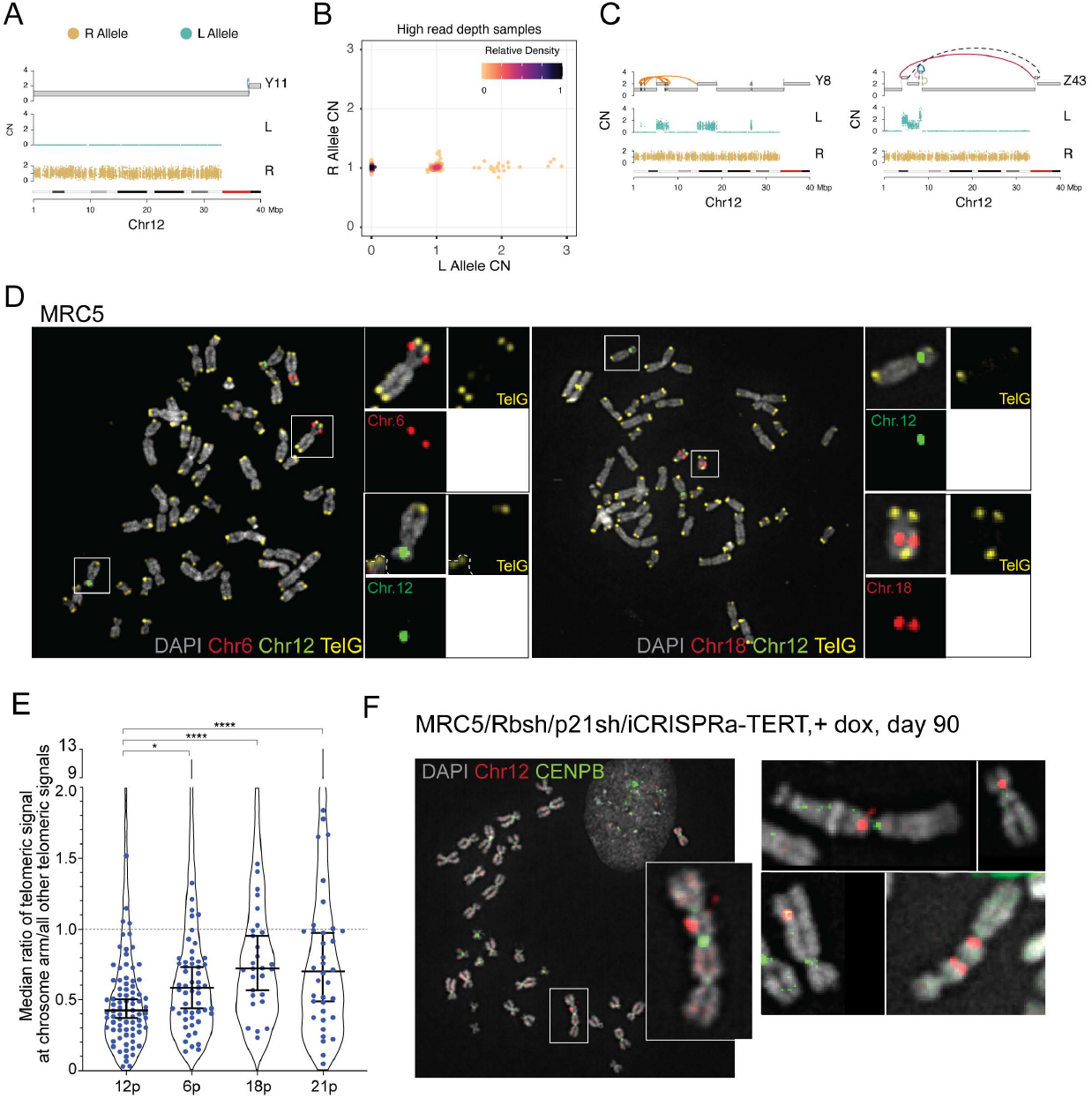
Telomere attrition renders an allele of 12p vulnerable. **A)** Genomic track plots of parental alleles phased into lost (“L”) and retained (“R”) haplotypes (see Methods) on chromosome 12p of clone Y11. **B)** Scatter plot showing purity- and ploidy-transformed L and R haplotype specific allelic read depth across 12p segments in high pass WGS-profiled post-crisis clones. **C)** Genomic track plots of allelic read counts on the L and R allele of clones Y8 and Z43, two post-crisis clones that independently acquired structural variants on an otherwise unrearranged chromosome 12p allele. **D)** Metaphase spreads of early passage MRC5 cells, hybridized with BAC probes to chromosome 12 (green) and either chromosome 6 or 18 (red), as well as a PNA probe for TelG (yellow). Insets of white-boxed chromosomes are shown with each channel individually. **E)** Quantification of the relative length of the shortest 12p telomere. Each dot shows the median ratio of the TelG signal of the shortest telomeres of the indicated chromosome arm to all other telomeres in each metaphase spread. Violin plots show data from all telomeres analyzed. Chromosome 12 was identified using a specific BAC probe (Chr.12p11.2) in 79 metaphase spreads with a total of 3992 telomeres. Chromosome 6 and 18 were identified based on BAC probe hybridization (Chr.6p21.2∼21.3, Chr.18q12.3∼21.1) in 53 and 28 metaphases, respectively (2629 and 1497 telomeres, respectively). Chromosome 21 was identified from DAPI banding patterns in 36 metaphases (1757 telomeres). *P* values were derived from an unpaired Students t-test, *, *p* <0.05, ****, *p* <0.0001. See Methods. **F)** Dicentric chromosomes containing chromosome 12 in telomere crisis. Metaphase spreads from MRC5/Rbsh/p21sh/iCRISPRa-TERT cells with doxycycline at day 90 (during crisis) were hybridized with a BAC probe for Chr.12p11.2 (red) and a CENPB PNA probe (green) to identify centromeres. A full spread is shown with white box inset zoom in. Further examples from other spreads are also shown. See also Supplementary Figure 7B.

We next tested whether the preferential 12p events could be due to the presence of a short telomere on one of the 12p alleles. To this end, we combined telomeric FISH with BAC probes specific for chromosome 12 and two other chromosome (6 and 8) that did not show evidence for structural variants in WGS (Figure 7D, Figure 4B). Comparing the ratio of the telomeric signal of the shortest 12p telomeres to the signal of all other telomeres in individual metaphase spreads revealed that one of the 12p telomeres was significantly shorter (Figure 7E). The shortest telomeres of 6 and 18 (Supplementary Figure 7B) were also shorter than the median but not to the same extent as 12p. The relative telomere length of the shortest 21p allele showed a heterogenous distribution that overall was significantly longer than 12p in the parental cells (Figure 7E). This does not exclude the possibility of 21 becoming critically short at later time points, and indeed the observation of a low percentage of clones in the 5X WGS screening with amplifications of 21q could indicate that this chromosome end did occasionally become deprotected in this population (Figure 4C). Such deprotection of a chromosome 21 telomere is consistent with chromosome 21 preferentially stabilizing the derivative chromosome 12 (Figure 6B, Supplementary Figure 6A).

To look for evidence of chromosome 12 being involved in the initial fusion events in this system we combined a chromosome 12 BAC probe with a centromere probe in MRC5/Rbsh/p21sh/iCRISPRa-TERT cells in crisis (at day 90). Strikingly, we observed a number of instances of chromosome 12 within chromosome fusion events (Figure 7F). The fraction of chromosome fusions involving chromosome 12 is higher than expected (∼50% observed versus ∼4% expected, Supplementary Figure 7C). Collectively, these data support the hypothesis that a short telomere on one allele of 12p increased the chance of 12p partaking in a fusion event that preceded subsequent rearrangement lineages.

## Discussion

We have described the first whole genome profiles of cells emerging from natural telomere crisis, both in the setting of spontaneous and controlled telomerase activation. Analysis of a variety of post-crisis genomes from divergent lineages and independent immortalization events uncovered highly complex patterns of copy number amplification and rearrangement. These genomes were not typified by the expected predominance of fold-back inversions that are indicative of BFB cycles or low amplitude copy number oscillations associated with chromothripsis. These cell lines spent a varying amount of time in telomere crisis, potentially with very different numbers of chromosome fusions, which is hard to quantify with limited historical data available. Although we consider it unlikely, we cannot rule out the possibility that some of the rearrangements we observed are not a direct consequence of telomere crisis. Due to the limited similarities between these cell lines, we constructed an *in vitro* system that allowed us to sequence high numbers of post-crisis genomes.

We consider our *in vitro* system to be a good representation of telomere crisis for a number of reasons. Telomeres in this system have been eroded through replicative attrition, rather than being subject to acute deprotection by the removal of TRF2. This is an important distinction since telomeres lacking TRF2 are repaired by c-NHEJ whereas other pathways are active at naturally eroded telomeres^35,38,39^. Furthermore, the number of dicentric chromosomes in our system is low (generally 1-2 per metaphase spread) which is similar to the frequency observed in other natural telomere crisis systems^29^. Apart from the abrogation of the Rb/p21 pathways, which is considered likely to occur before telomere crisis *in vivo*^40–43^, these cells contain intact DNA repair pathways, and we make no assumptions as to what the predominant repair mechanisms will be in this context. Further, the relatively weak telomerase activity that can only sustain the shortest telomeres within the population is similar to what occurs in cancer, since many tumours maintain very short telomeres despite activation of telomerase^44–47^.

This system revealed striking convergent evolution of rearrangements on chromosome 12p, for which we consider the most plausible explanation to be a short telomere on one of the 12p alleles driving the first chromosome fusions in telomere crisis. Although the rearrangement events on 12p are likely specific to this cell line, we can draw valid conclusions about the consequence of short telomeres across other systems. It seems likely that the first events during telomere crisis are driven by the shortest telomere(s) within a cell population. We define a minimal set of events that can occur as a result of a single deprotected telomere. We document clean patterns of BFB-like events that represent progressive stages in the evolution of more complex genome architectures. This data has provided an important snapshot into the events that occur during a relatively short time period of telomere crisis. The comparatively flat genomes in the majority of post-crisis clones suggest that the consequences of telomere crisis do not have to be spectacular. It may be that in this system there is selection against complex events involving multiple chromosomes. The more complex events that can be observed in the immediate aftermath of dicentric chromosome resolution^14^ may not lead to viable post-crisis clones. Our data also point to a surprising role for acrocentric chromosomes in stabilizing fusion events, which has also been suggested and observed in other studies^14,48^. Since it was technically challenging to resolve these stabilizing events using solely WGS analysis, it is possible that these types of events have been overlooked in large-scale WGS analyses and could be an important hallmark of post-crisis genomes.

In summary, our data reveal that telomere crisis can instigate a wide spectrum of structural variations in the viable descendants of this genomic trauma. Since no single type of variation appears to be a hallmark of past telomere crisis, other genomic insignia of telomere crisis will have to be identified in order to determine whether a given cancer has experienced telomere dysfunction in its proliferative history.

## Methods

### Cell lines

MRC5 human lung fibroblasts (CCL-171), Phoenix-ampho (CRL-3213), RPE-1 hTERT (CRL-4000), HCT-116 (CCL-247) and U2-OS (HTB-96) cells were obtained from ATCC for this study. 293-FT cells were obtained from ThermoFisher. MRC5 cells and derivatives thereof were grown in EMEM media (ATCC) supplemented with 15% fetal bovine serum (FBS; Gibco) and 100 U/mL of penicillin and 100 μg/mL streptomycin (PenStrep, Gibco) at 37°C, 5% CO_2_. hTERT RPE-1 cells were grown in DMEM:F12 media (Gibco) with 10% FBS and PenStrep at 37°C, 5% CO_2_. HCT-116 colorectal carcinoma cells and U2OS cells were grown in DMEM with 10% FBS and PenStrep at 37°C, 5% CO_2_.

### Immortalized cell line panel

Details of the post-crisis immortalized cell line panel are provided in Supplementary Table 1. HA-1M cells were a kind gift of Silvia Bacchetti^29^, SW13/26/39 cells were a kind gift of Jerry Shay^28^, and Bet-3B/3K and BFT-3B/G/K cells were a kind gift of Roger Reddel^27^.

### Cloning and plasmids

A dual-shRNA vector LM2PshRB.698-p21.890-PURO was used to knockdown Rb and p21^49^. The inducible dCas9-VPR (pCW57-dCas9-VPR) construct was created by Gibson assembly of the dCas9-VPR insert from SP-dCas9-VPR (Addgene#63798) into pCW57-MCS1-P2A-MCS2-Neo (Addgene#89180). Retroviral pLVX-hTERT was a kind gift of Teresa Davoli. Activating *TERT* gRNAs were targeted up to 1000 bp upstream of the *TERT* promoter transcriptional start site, and designed using online software from the Broad Institute (portals.broadinstitute.org/gpp/public/analysis-tools/sgrna-design). gRNA sequences were cloned into a modified version of lentiGuide-Puro (Addgene#52963) in which the selection cassette had been swapped for Zeocin resistance. Activating *TERT* gRNA sequences are shown in Supplementary Table 3. *TTN* gRNA sequences were used as described^32^.

### Viral gene delivery

Retroviral constructs were transfected into Phoenix amphitropic cells using calcium phosphate precipitation. Lentiviral constructs were transfected with appropriate packaging vectors using calcium phosphate precipitation into 293-FT cells. Viral supernatants were collected and filtered before addition to target cells, supplemented with 4 μg/ml polybrene. For activating gRNA constructs, multiple viral supernatants were collected and concentrated using PEG-it Virus Precipitation Solution (System Biosciences LV810A-1). Cells were infected 2-3 times at 12-hour intervals before selection in the appropriate antibiotic.

### Immunoblotting

For immunoblotting, cell pellets were directly lysed in 1X Laemmli buffer (2% SDS, 5% *β*-mercaptoethanol, 10% glycerol, 0.002% bromophenol blue and 62.5 mM Tris-HCl pH 6.8) at a concentration of 10^7^ cells/ml. Lysates were denatured at 100°C, and DNA was sheared with a 281/2 gauge insulin needle. Lysates were resolved on SDS/PAGE gels (Life Technologies), transferred to nitrocellulose membranes and blocked with 5% milk in TBS with 0.1% Tween-20. Primary antibodies (anti-Cas9 7A9-3A3, Cell Signaling Technology #14697S, anti-γ-tubulin Sigma#T5326, anti-Human Retinoblastoma protein BD Pharmigen #554136, anti-p21 F-5 Santa Cruz sc-6246) were incubated overnight, before membrane washing and incubation with appropriate HRP-conjugated secondary antibodies (Amersham) and detection with SuperSignal ECL West Pico PLUS chemiluminescence (ThermoFisher).

### Immunofluorescence

Cells were grown on glass coverslips and fixed in 3% paraformaldehyde and 2% sucrose. Coverslips were permeabilized in 0.5% Triton-X-100/PBS, and blocked in goat block (0.1% BSA, 3% goat serum, 0.1% Triton-X-100, 2mM EDTA) in PBS. Primary and secondary antibodies (Rabbit anti-53BP1 Abcam #ab-175933, F(ab’)2-Goat anti-Rabbit IgG (H+L) Cross-Adsorbed Alexa Fluor 488 ThermoFisher A-11070) were diluted in goat block. Slides were counter-stained with DAPI and mounted using Prolong gold anti-fade medium. Images were acquired on a DeltaVision microscope (Applied Precision) equipped with a cooled charge-coupled device camera (DV Elite CMOS Camera), with a PlanApo 60× 1.42 NA objective (Olympus America), and SoftWoRx software. Images were analyzed for foci numbers using a custom-made algorithm written for FIJI, courtesy of Leonid Timashev^50^.

### Metaphase spread preparation and staining

Metaphase spreads were prepared by treatment of cells with 0.1 ug/ml colcemid (Roche) for 3 hours, before trypsinization and swelling at 37°C for 5-10 mins in 0.075 M KCl. Cells were fixed in a freshly prepared 3:1 mixture of methanol to acetic acid and stored at 4°C at least overnight. Spreads were prepared by dropping cell solution onto cold glass slides exposed to steam from a 75°C water bath, flooding slide with acetic acid, before exposure of the dropped cells for 3-5 secs in steam. Slides were dried overnight before storage in 100% Ethanol at -20°C. For visualization of fusions, slides were rinsed in PBS, fixed in 4% formaldehyde/PBS for 5 mins, and dehydrated in an ethanol series before co-denaturation of slide and PNA probes (TelG-Cy3 PNA Bio F1006, CENPB-AF488 PNA Bio F3004) for 3 mins at 80°C in hybridization solution (10 mM Tris-HCl pH 7.2, 70% formamide, 0.5% Roche 11096176001 blocking reagent). Hybridization was carried out for 2 hours at RT in the dark, before washing twice in 10 mM Tris-HCl pH 7.2, 70% formamide and 0.1% BSA, then washing three times in 0.1 M Tris-HCl pH 7.2, 0.15M NaCl, 0.08% Tween-20. DAPI was included in the second wash. Slides were dehydrated through an ethanol series before mounting with Prolong Gold antifade medium (Invitrogen).

For chromosome painting, slides were prepared as above for chromosome fusions, and co-denaturation of chromosome specific paints (XCP-12 Metasystems D-0312-050-FI, XCP-21 Metasystems D-0321-050-OR) was carried out at 75°C for 2 mins, before hybridization overnight at 37°C. Post hybridization washes were 0.4X SSC for 2 mins at 72°C, 2X SSC, 0.05% Tween-20 for 30 secs, followed by counterstaining in DAPI for 15 min, and a rinse in ddH_2_O before mounting in Prolong Gold antifade medium (Invitrogen). For karyotyping, slides were prepared as above, and analysis was carried out on DAPI stained chromosomes.

### BAC probes

To identify individual chromosomes on metaphase spreads, BAC probes were ordered from BACPAC Genomics (Chr.12p11.2 RP11-90H7, Chr.18q12.3∼21.1 RP11-91K12, Chr.6p21.2∼21.3 RP11-79J17). Probe DNA was nick-translated with either Digoxigenin-11-UTP or Biotin-16-UTP (Roche) using DNase I (Roche) and DNA polymerase I (NEB) overnight at 15°C. Probes were precipitated with Cot1 Human DNA (Invitrogen) and salmon sperm DNA (Invitrogen) and resuspended in 50% formamide, 2X SSC and 10% dextran sulfate before denaturation for 8 mins at 80°C. Metaphase spreads were prepared as above, and slides were denatured with 70% formamide, 2X SSC for 2 mins at 80°C before dehydration through an ethanol series. Slides were co-denatured for 2 mins at 80°C with TelG-647 (PNA Bio F1014) in hybridization solution (10mM Tris-HCl pH 7.2, 70% formamide, 0.5% Roche 11096176001 blocking reagent) followed by a 2-hour hybridization at RT. Denatured BAC probes were then applied and hybridized overnight at 37°C. Slides were washed for 3 x 5 mins in 1X SSC at 60°C, followed by a blocking step in 30 μg/ml BSA, 4X SSC and 0.1% Tween-20 for 30 mins at 37°C. BAC probes were detected with anti-Digoxenin-Rhodamine (Roche 11207750910) and Avadin-FITC antibodies (VWR CAP21221) in 10 μg/ml BSA, 1X SSC and 0.1% Tween-20 by incubating for 30 mins at 37°C, before washing twice for 5 mins in 4X SSC and 0.1% Tween-20 at 42°C.

Counterstaining with DAPI was carried out for 15 mins at RT, before a further wash at 42°C in 4X SSC, 0.1% Tween-20 and mounting in Prolong-Gold antifade (Invitrogen). Images were acquired on the Deltavision microscope equipment detailed above. Images were analyzed using FIJI^51^. Briefly, individual chromosomes were detected with the BAC probes. Intensity measurements of the Tel-G signal were quantified for each p-arm and q-arm of identified chromosomes. Measurements were also taken for all other telomeres in the same spread. Background subtracted measurements for all telomeres were compared to the shortest (lowest intensity) of the Chr.12p (or Chr.18p or Chr.6p) telomeres on each spread, and the results expressed as a ratio. For identification of fusions containing chromosome 12, the same protocol was carried out using a centromere probe (CENPB-AF488, PNA Bio F3004).

### qPCR

RNA was isolated from cell pellets using a Qiagen RNeasy kit, according to the manufacturer’s instructions. cDNA was synthesized using Superscript IV first-strand synthesis (ThermoFisher). qPCR was carried with SYBER Green reagents (ThermoFisher) and run on a Life Technologies QuantStudio 12K machine. qPCR primer sequences are shown in Supplementary Table 3.

Expression was quantified using the standard *Δ*CT method relative to *β*-actin.

### TRAP assay

Telomerase activity was assessed using the TRAPeze kit (EMD Millipore S7700) according to the manufacturer’s instructions. Amplification products were resolved on 12% PAGE gels and visualized with EtBr staining.

### STELA, Fusion PCR and telomeric blots

High molecular weight DNA was extracted from cell pellets using a MagAttract HMW DNA kit (Qiagen) and solubilized by overnight digestion with *EcoR*I (for STELA and Fusion PCR) or a combination of *Alu*I and *Mbo*I (for telomeric blots). STELA was carried out essentially as described^34^. Briefly, 10 ng of DNA was annealed to a mixture of six telorette linkers (Supplementary Table 3) overnight at 35°C, before dilution with water to a concentration of 200 pg/μl. Multiple PCR reactions for each sample were carried out with 200 pg of annealed DNA using the XpYpE2 and teltail primers (Supplementary Table 3) and FailSafe PCR reagents (Epicentre). PCR conditions were as follows: 94°C for 15s, 27 cycles of 95°C for 15s, 58°C for 20s, 68°C for 10 mins, and a final extension at 68°C for 9 mins. PCR products were resolved on 0.8% TAE gels, denatured and transferred to Hybond membrane via Southern blotting. Products were detected with a randomly primed *α*-^32^P DNA probe created by amplification of the telomere-adjacent region of the XpYp telomere (using XpYpE2 and XpYpB2 primers, Supplementary Table 3). For quantification, FIJI was used to measure relative signal between indicated molecular weight markers relative to background signal for each sample.

Fusion PCR was carried out essentially as described^11,35^. Subtelomeric primers (Supplementary Table 3) used for amplification of telomeric fusions were XpYpM, 17p6 and 21q4. The control primer XpYpc2tr was included for control amplification of XpYp subtelomeric DNA and detected using EtBr. Fusion products were detected with a random primed *α*-^32^P labelled DNA probe (21q probe) specific for the TelBam11 telomere subfamily^36,52^, which was created with the 21q4 primer and 21q-seq-rev2. The number of fusions per haploid genome (6 pg) is calculated based on the amount of input DNA in each PCR reaction.

Telomere length was assessed using telomeric restriction fragment analysis. Briefly, *Alu*I/*Mbo*I-digested genomic DNA was run on 0.8% TAE gels, before denaturation, neutralization, and transfer onto a Hybond membrane according to standard Southern blotting procedures. Telomeric DNA was detected using a *α*-^32^P labelled Sty11 telomeric repeat probe^45^.

### WGS Library preparation

Genomic DNA was extracted from cell pellets using a QIAGEN QIAamp DNA mini kit and sheared using a Covaris Ultrasonicator (E220) to approximately 300 bp fragments. DNA concentration was measured using Qbit 4.0 reagents (ThermoFisher) and 200 ng of fragmented DNA was used for library preparation. End repair and A-tailing was carried out with NEBNext End repair reaction enzyme mix and buffer (E7442), and KAPA Dual-Indexed Adapters (Roche) were ligated using the T4 DNA ligase kit from NEB (M0202). Post-ligation size selection was performed with AMPure XP beads (Beckman Coulter) before washing two times in 80% ethanol. Libraries were amplified using KAPA HiFi HotStart ready mix (Roche) and P5 and P7 primers (IDT). PCR program was as follows: 98°C for 45 s, 5 cycles of 98°C for 15 s, 60°C for 30 s, 72°C for 30 s and a final extension at 72°C for 5 mins. A further size selection and washing step was carried out after library amplification, and library quality was confirmed on Bioanalyzer chips (Agilent) and using a KAPA Library Quantification kit (Roche). Libraries were pooled and submitted for sequencing on NovaSeq 6000 at the New York Genome Center.

### WGS basic data processing

Reads were aligned to GRCh37/hg19 using the Burroughs-Wheeler aligner (bwa mem v0.7.8, http://bio-bwa.sourceforge.net/)^53^. Best practices for post-alignment data processing were followed through use of Picard (https://broadinstitute.github.io/picard/) tools to mark duplicates, the GATK (v.2.7.4) (https://software.broadinstitute.org/gatk/) IndelRealigner module, and GATK base quality recalibration.

Variant rearrangement junctions were identified using SvAbA^54^ (https://github.com/walaj/svaba) and GRIDSS^31^ (https://github.com/PapenfussLab/gridss) with standard settings. For MRC5 samples, the somatic variant setting of each tool was used, with the ancestral MRC5 line as the matched normal. SvAbA was applied using a panel of normals (PON) that was constructed by running SvABA to obtain constitutional junction calls for 1,017 TCGA tumor/normal pairs (TCGA dbGaP: phs000178.v11.p8). For GRIDSS, a PON was obtained from the Hartwig Medical Foundation (https://nextcloud.hartwigmedicalfoundation.nl). 1 kbp binned GC and mappability corrected read depth was computed using fragCounter (https://github.com/mskilab/fragcounter). Systematic read depth bias was subsequently removed using dryclean (https://github.com/mskilab/dryclean)^55^.

### Low-pass WGS clustering

Genome-wide binned read depth was aggregated across 118 low pass WGS clones across 10 kbp bins by taking the median of 1 kbp binned normalized read depth from dryclean (see above). To minimize read depth noise in unmappable regions, recurrent (>10% of the cohort) low-quality coverage regions (defined in ref.^3^) are combined with regions bearing consistently high variance in our high-pass sequencing dataset (standard deviation >0.3 for bin value over the mean in 100 kbp windows). Hierarchical clustering was then applied on the genome-wide Euclidean distance of bins, with “method = ward.D2” option. Six clusters were identified following dendrogram inspection.

### Junction balance analysis

Preliminary junction balanced genome graphs were generated for MRC5 and SV40T cell lines from binned read depth and junction calls (see above) using JaBbA (https://github.com/mskilab/JaBbA)^3^. Briefly, 1 Kbp binned read depth output from dryclean was collapsed to 5 Kbp and JaBbA was run with slack penalty 500. gGnome (https://github.com/mskilab/gGnome) was used to identify complex structural variant patterns. Genome graphs and corresponding genomic data (e.g. binned coverage, allelic bin counts) were visualized using gTrack (https://github.com/mskilab/gTrack).

### Joint inference of junction balance in MRC5

To chart structural variant evolution across sub-clades of MRC5 clones, a procedure was developed to jointly infer junction balanced genome graphs in a lineage (e.g. BFB lineage in Figure 5C). This co-calling algorithm augmented the existing JaBbA model, described in detail in^3^, enabling application to a compendium of genome graphs by minimizing the total number of unique loose ends assigned a nonzero copy number across the graph compendium.

To describe this algorithm, we extend the notation introduced in ref.^3^. Formally, we define a collection {*G*^*i*^}_*i*∈1,…*n*_of identical genome graphs across clones, each a replica of a “prototype” genome graph *G*^*0*^. The mapping *p* maps each vertex *ν* ∈ *V*(*G*^*i*^) and edge *e* ∈ *E*(*G*^*i*^), *i* ∈ 1, …*n* to its corresponding vertex *p*(*ν*) ∈ *V*(*G*^0^) and edge *p*(*ν*) ∈ *V*(*G*^0^) in the prototype graph. We then jointly infer unique copy number assignments *k*^*i*^ to the vertices and edges of each genome graph *G*^*i*^ by solving the mixed integer program:

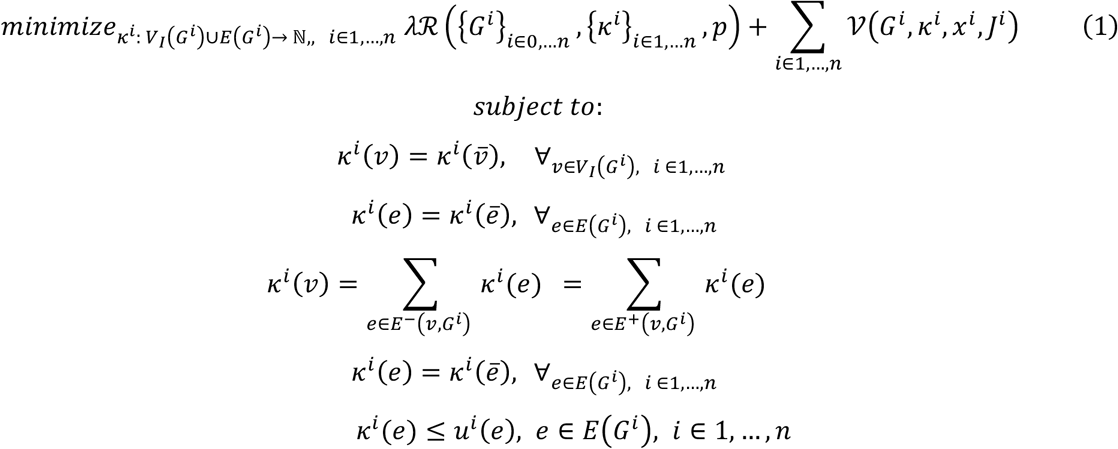

where *x*^*i*^ and *J*^*i*^ represent the binned read depth data and bin-node mappings for clone *i* and 𝒱(*G*^*i*^, *k*^*i*^, *x*^*i*^, *J*^*i*^) is the read depth residual for genome graph *i*, analogous to^3^. An addition term in this new joint formulation is *μ*^*i*^: *E*(*G*^*i*^)→{0,∞}, which is a data derived mapping that constrains the upper bound of each edge *e* ∈ *E*(*G*^*i*^), e.g. on the basis of whether that junction has read support in clone *i*. In addition, a joint complexity penalty *R* couples the collection *k*^i^}of copy number mappings across the collection of graphs {*G*^*i*^}_*i*∈1,…*n*_to each other by jointly penalizing loose ends at all vertices that map to the same prototype graph vertex *ν* ∈ *V*(*G*^0^). Formally,

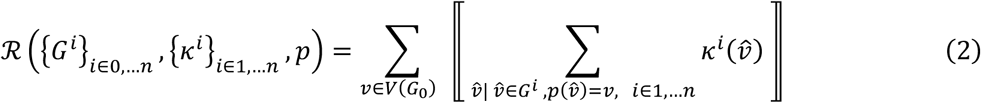

As in ^3^, the hyperparameter λ in Equation 1 controls the relative contribution of the read-depth residual and complexity penalty to the objective function. It is important to note that while each of the graphs *G*^*i*^ have an identical structure, the constraints imposed by the upper bounds *u*^*i*^ and bin profiles *x*^*i*^ couple each graph to its junction and read depth data, and hence lead to a unique fit *k*^*i*^ on the basis of this data. The ℓ_*0*_penalty (defined using the Iversion bracket ⟦ ⟧operator) in Equation 2 couples the solutions *k*^*i*^ by adding an exponential prior on the number of unique loose ends across entire graph compendium, where uniqueness is defined by the mapping *p* to the prototype graph *G*^0^.

This joint mixed-integer programming model in Equation 1 is implemented in the “balance” function of gGnome. The model was applied to a collection of genome graphs representing the structure of chromosome 12 across 13 clones. The prototype graph for this genome graph collection was built from the disjoint union of intervals of the 13 preliminary graphs (via the GenomicRanges “disjoin” function) and the union of junction calls fit across those graphs (via gGnome “merge.Junction” function). Each graph was associated using the read depth data and bin-to-node mappings as per ^3^. The mapping *u*^*i*^ for each reference edge was set to ∞ while variant edges were assigned ∞ on the basis of bwa mem realignments of read pairs in each clone .bam file to the corresponding junction contig via rSeqLib (https://github.com/mskilab/rSeqLib)^56^, otherwise they were assigned 0.

Equation 1 was then solved using the IBM CPLEX (v12.6.2) MIQP optimizer within the gGnome package after setting the hyperparameter λ to 100. This value was chosen after a parameter sweep observing for the visual concordance of genome graphs, loose ends, and read depth profiles in the region.

### Joint reconstruction of allelic evolution in MRC5

Evolving 12p alleles were jointly reconstructed across 13 MRC5 clones through the analysis of junction balanced genome graphs (*G*^*i*^,*k*^*i*^) (see “Joint inference of junction balance in MRC5” section above). The procedure for joint allelic phasing described in ^3^ was extended to identify the most parsimonious collection of linear and/or cyclic walks and associated walk copy numbers that summed to the vertex and edge copy numbers in the compendium (*G*^*i*^, *k*^*i*^).

Formally, the subgraph of vertices and edges with a nonzero copy number in each (*G*^*i*^,*k*^*i*^) were exhaustively traversed to derive all minimal paths and cycles *H*^*i*^, where for each walk *h* ∈ *H*^*i*^ maps to subsets *V*(*h*)⊆*V*(*G*^*i*^) and *E*(*h*)⊆*E*(*G*^*i*^) of vertices and edges in the graph *G*^*i*^. The nodes and vertices of these walks were then projected to via the mapping *p* to define a unique set of walks *H*^0^ in the prototype graph *G*^0^. We extend our notation *p* (see previous section) so that for a walk *h* ∈ *H*^*i*^ the mapping *p*(*h*) ∈ *H*^0^ denotes the walk formed by projecting the vertices and edges of *h* via *p* to *H*^0^. With these definitions, the single graph haplotype inference defined in ^3^ was extended to a joint inference by solving the following mixed integer linear program to assign a copy number *ϕ*^*i*^(*h*) ∈ ℕ to each walk *h* ∈ *H*^*i*^.

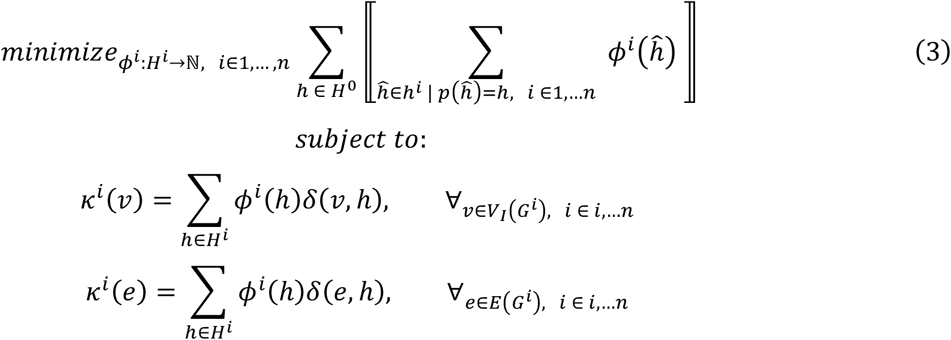

where the function *δ*(*ν, h*) and *δ*(*e,h*) is 1 if vertex *ν* and edge *e* belong to walk *h* and 0 otherwise. The Iverson bracket (⟦ ⟧) operator in the objective function Equation 3 minimizes the total number of unique walks used across the compendium, hence identifying a jointly parsimonious assignment of copy number to walks across the compendium of graphs. Equation 3 was solved using the IBM CPLEX (v12.6.2) MIQP optimizer within the gGnome package. Variant cycles and paths from the resulting solution were manually combined to yield a set of consistent linear paths, i.e. somatic haplotypes, to yield allelic reconstructions in Figure 5C.

### Loose end classification

Each loose end in each MRC5 genome graph was analyzed to identify a clone specific (i.e. absent in the ancestral MRC5 line) origin for the mates of high mapping quality (MAPQ=60) reads mapping to the location and strand of the loose end. These mates were assessed for neo-telomeric sequences by counting instances of 11 permutations of a 12-bp telomere repeat motif (TTAGGGTTAGGG) (using the R / Bioconductor Biostrings package) in the mates. The mates were also assembled into contigs using fermi^57^ aligned using bwa mem^58^ via the RSeqLib R package^56^ to hg38 (which contains a more highly resolved centromere build) to characterize novel repeat (e.g. centromere) fusions (GATK Human reference genome, hg38, data bundle including Homo_sapiens_assembly38.fasta, gs://gcp-public-data--broad-reference). The loose end loci were also assessed through overlap with the hg19 repeatMasker database (human_g1k_v37_decoy.repeatmasker) for the presence of reference annotated repeats that might explain the absence of a mappable junction explaining the copy number change.

### SNV phylogeny

To compute an SNV phylogeny across MRC clones, we first identified SNV that were acquired in MRC5 clones relative to the ancestral MRC5 line using Strelka2^59^ (https://github.com/Illumina/strelka) under paired (i.e. tumor / normal) mode with the clone as the “tumor” and the MRC ancestral line as the “normal” sample and default parameters and GATK hg19 resource bundle (Genome Analysis Toolkit GATK Resource Bundle for hg19; gs://gatk-legacy-bundles). Acquired SNVs were first filtered according to the Strelka2 PASS filter as well as additional filters (MQ = 60, SomaticEVS>12, total ALT count > 4) yielding 27,220 total unique variants across the 13 MRC5 clones. Reference and variant allelic read counts were assessed at each SNV site (via the R / Bioconductor Rsamtools package, version 3.6.1, http://www.r-project.org/) across all 13 clones. We then further required a >0.5 posterior probability of a variant being present in a sample, by assuming Binomial likelihood of variant read count and using the aggregated allele frequency in all samples as the prior, resulting in the final 14,970 unique mutations. The binary matrix of clones by SNV loci. was then used to derive a neighbor-joining phylogenetic tree using the R / Bioconductor package ape. Following tree construction, we associated each SNV with its most likely phylogenetic tree branch by comparing the binary incidence vector associated with each SNV with the binary incidence vector associated with each tree branch, and finding the closest branch using Jaccard distance, only linking SNV to branches when the SNV was within <0.1 Jaccard distance of the closest branch, thus producing the groupings of SNVs in Figure 5A.

### Parental SNP allelic phasing and imbalance

Germline heterozygous sites in the parental MRC5 line were identified by computing allelic counts at HapMap sites (GATK human reference genome, hg19 data bundle, hapmap_3.3.b37.vcf) and identifying loci with variant allele fraction >0.3 and <0.7. Y11, a clone with loss of a single allele at 12p, was chosen to phase parental SNPs on 12p. At each locus, the allele (reference or alternate) with a 0 read count was assigned to the “L” (lost) haplotype and the other allele was assigned to the “R” (retained) haplotype. (All heterozygous SNP loci in the region contained exactly one allele with a 0 read count). L and R allelic counts were then computed at these sites across all 13 high pass WGS and 131 low pass WGS samples. These counts were divided by the genome wide mean of heterozygous SNP allele counts (in these 100% pure and nearly diploid samples) to derive the absolute allelic copy number^60^.

### SNV clustering

Inter-SNV distances were computed for all pairs of reference adjacent acquired SNVs associated with each MRC5 clone and visualized as rainfall plots. Runs of two or more SNVs with inter SNV distances < 2 Kbp were nominated as clusters. Two distinct SNV clusters were identified on chromosome 12p across the 13 clones.

### Statistical Analysis

Statistical analysis for in vitro experiments was carried out using Prism software (GraphPad Software). All relevant statistical experimental details (n numbers, SD) are provided in the figure legends. Statistically significant associations between binary variables were determined using two-tailed Fisher’s exact test. Significance was assessed on the basis of Bonferroni-corrected *P* values < 0.05. Effect sizes (odds ratios) are reported alongside 95% confidence intervals for each test. Details of all other quantitative analyses (e.g. read depth processing, clone clustering, genome graph inference, allelic reconstructions, phylogenetic reconstruction, SNV clustering, parental SNP phasing) are described above.

## Data and Code Availibility

Custom software packages referenced in this study are available at https://github.com/mskilab (JaBbA build d7f4bff, gGnome build c998026, dryclean build 6d2bced, fragCounter build 575af99, rSeqLib build 23fbaf0, skitools build 61187fa, gUtils build 449ab2a, gTrack build 947c35c). Analysis code to generate the figures in the paper is available on request. Whole genome sequencing data has been deposited to the sequence read archive as aligned .bam files (https://www.ncbi.nlm.nih.gov/sra) under the identifier SUB8078633 (submission pending).

## Acknowledgements

We thank Jerry Shay (UTSW), Silvia Bacchetti, AdVec and Roger Reddel (CMRI) for the generous provision of cell lines used in this study. S.M.D is funded jointly by a NIH/NCI grant (5R35CA210036) and Melanoma Research Alliance grant (577521) both awarded to T.d.L. T.d.L has additional funding support from NIH/NIA (5R01AG016642-21A1), Starr Cancer Consortium (I13-0019), Breast Cancer Research Foundation and a Glenn Foundation Award from the Glenn Foundation. M.I., X.Y., J.B., and H.T. are supported by Burroughs Wellcome Fund Career Award for Medical Scientists, Doris Duke Clinical Foundation Clinical Scientist Development Award, Starr Cancer Consortium Award, and National Institutes of Health (NIH) U24-CA15020 to M.I., as well as Weill Cornell Medicine Department of Pathology Laboratory Medicine startup funds.

## Author contributions

Conceptualization: S.M.D, M.I, T.d.L

Methodology: S.M.D, Y.X, M.I

Software and Formal Analysis: Y.X, J.R, J.B, M.I

Investigation: S.M.D, N.B, K.T, H.T

Writing – Original Draft: S.M.D, M.I, T.d.L

Writing – Review and Editing: S.M.D, Y.X, M.I, T.d.L

Supervision and Funding Acquisition: T.d.L, M.I

## Supplementary Figure Legends

**Supplementary Figure 1. Complex gains and telomerase restoration after spontaneous telomere crisis resolution. Related to Figure 1**.

**A)** TRAP assay showing telomerase activity in SV40 immortalized clones SW13, SW26 and SW39. MRC5 is included as a negative control, and HCT116 as a positive control. TSR8 is the positive control template.

**B-C)** Example clusters of complex gains across the cell lines shown in Figure 1A, each showing binned purity- and ploidy-transformed read-depth, with the top track showing the associated junction-balanced genome graph (see Methods, ^3^, with y-axis representing units of per cell copy number (CN) across bins and graph nodes (i.e. intervals). Gray and colored edges represent reference and variant junctions, respectively. Blue edges represent loose ends (see Methods for further details). Bins and junctions are colored as per Figure 1A.

**Supplementary Figure 2. Controlled system for crisis escape. Related to Figure 2**.

**A)** Immunoblot for Rb and 21 in MRC5 cells and MRC5/Rbsh/p21sh cells.

**B)** Schematic diagram illustrating the iCRISPRa-TERT system. The positions of the *TERT* activating gRNAs relative to the transcriptional start site (TSS) of the *TERT* gene are shown.

**C)** qPCR for *TTN* control gene, activated with a combination of four sgRNAs in MRC5/Rbsh/p21sh iCRISPRa cells (with or without dox, 96 hrs). HCT116, RPE1-hTERT and U2OS cells are included as negative controls. Data are from three independent biological replicates.

**D)** STELA of the XpYp telomere in MRC5/Rbsh/p21sh/iCRISPRa-TERT cells after 70 days or 150 days of continuous culture with or without doxycycline. This is a biological replicate of the STELA in Figure 2E.

**Supplementary Figure 3. Telomere fusions after controlled escape from crisis. Related to Figure 3**.

**A)** Quantification of percentage of fused chromosomes after the indicated days of continuous culture for MRC5/Rbsh/p21sh/iCRISPRa-TERT cells with and without doxycycline. Data represent means and SDs for three independent biological replicates. *P* values derived from an unpaired Student’s t-test. ns, not significant; **, *p* <0.01; ***; *p* <0.001.

**Supplementary Figure 4. Post-crisis telomere dynamics. Related to Figure 4**.

**A)** Products of telomere fusion PCR on a panel of post-crisis clones (See Figure 4A). Telomere fusions are detected by hybridization to the 21q probe. The control XpYp band is detected with Ethidium bromide staining.

**B)** STELA products from a panel of post-crisis clones from both time points.

**C)** Telomeric blot on DNA from MRC5 cells and a panel of post-crisis clones from both time points.

**D)** TRAP assay showing telomerase activity in MRC5/Rbsh/p21sh/iCRISPRa-TERT treated with doxycycline for the indicated number of days and a selection of post-crisis clones from the day 150 timepoint. HCT116 is included as a positive control.

**E)** TRAP assay showing robust telomerase activity in MRC5 cells infected with retroviral pLVX-hTERT.

**F)** Circular heatmap showing genome-wide binned purity- and ploidy-transformed read depth (in units of CN across 8 low pass WGS-profiled control CT clones). Heatmap rows correspond to concentric rings in the heatmap.

**Supplementary Figure 5. WGS analysis showing genome alterations post crisis clones. Related to Figure 4 and 5**.

**A)** Consensus copy number profile of 47 complex low pass WGS clones targeting chromosome 12p.

**B)** Rainfall plot demonstrating SNV patterns as a function of GC vs AT reference nucleotide context, with y-axis showing the logarithm of the inter SNV distance. Highlighted region represents a GC strand coordinated cluster that is found across all four clones in the BFB cluster which harbor junction j2.

**C)** Junction supporting read pairs and corresponding drop in coverage at the small deletion on chromosome 12q in representative BFB-like (Z43), chromothripsis-like (Y8), close-relative Y15, and lack of such evidence in distant Y11.

**D)** Detailed JaBbA models and linear allele reconstruction at two distinct loose ends, e2 and e3.

**Supplementary Figure 6. Karyotype evolution in post-crisis clones. Related to Figure 6**.

**A)** DAPI banded karyotypes from the MRC5 parental cell line and a selection of post-crisis clones. Marker chromosomes (chromosome 12) are indicated with a red star. Dashed box indicates the absence of an intact copy of chromosome 21.

**B)** Representative images of a metaphase spreads from clones Z41 and Z29 hybridized with both 12 (green) or 21 (red) chromosome paints. DNA stained with DAPI (gray).

**Supplementary Figure 7. Chromosome 12p allelic imbalance. Related to Figure 7**.

**A)** Scatter plot showing purity- and ploidy-transformed L and R haplotype specific allelic read depth across 12p segments in low pass WGS-profiled post-crisis clones.

**B)** Analysis showing that 12p, 6p and 18p arms contain the shortest telomere of those chromosomes. Comparison of the shortest (S) telomere end from each arm (p or q) to each other allele of that chromosome (i.e. q-short allele vs. p-short allele) across chromosome 12, 6 and 18 alleles. Ratio of telomeric intensity from TelG hybridization, as in Figure 7D (S= short allele, L= long allele, based on TelG intensity). Error bars show median and interquartile ranges. Chromosome 12: n= 49 cells, chr.18: n=27, chr.6: n= 22.

**C)** Fraction of dicentric or multicentric chromosomes containing 12p in MRC5/Rbsh/p21sh/iCRISPRa-TERT cells in crisis (with doxycycline at day 90) among all dicentric and multicentric chromosomes. Based on analysis of images as in Figure 7E. n=81 metaphases scored from two independent experiments.

## Supplementary Table Legends

**Supplementary Table 1: SV40T post-crisis cell lines. Related to Figure 1**.

Description of the SV40T immortalized cell lines used in this study (See Figure 1). Details of parental cell line, telomerase status and references are shown.

