## Supplemental Information for "Structural variant evolution after telomere crisis"

Dewhurst, Yao *et al.* Supplementary Figure 1

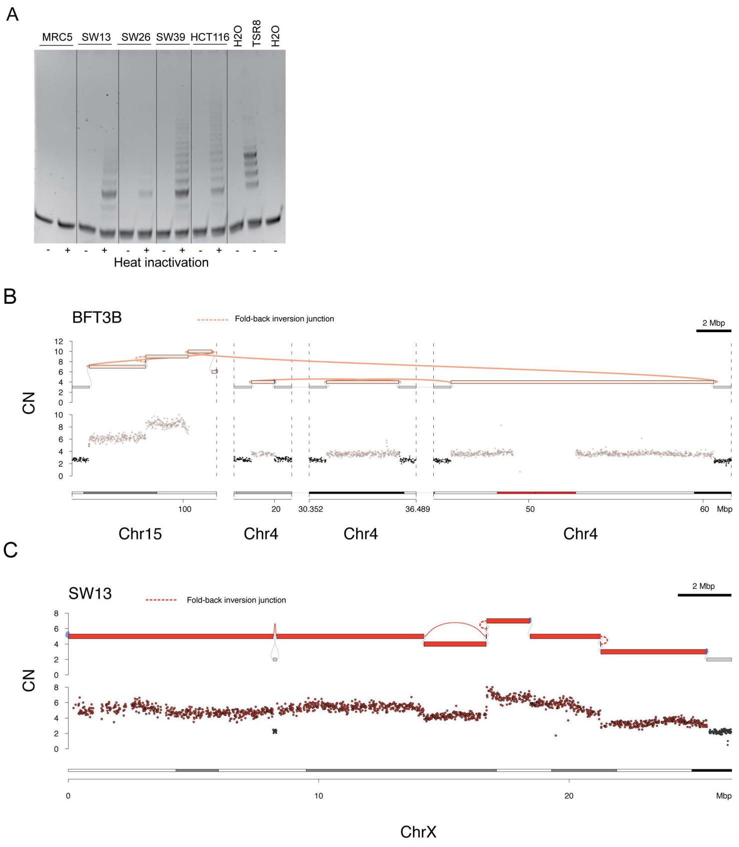


**Supplementary Figure 1. Complex gains and telomerase restoration after spontaneous telomere crisis resolution. Related to Figure 1.**

**A)** TRAP assay showing telomerase activity in SV40 immortalized clones SW13, SW26 and SW39. MRC5 is included as a negative control, and HCT116 as a positive control. TSR8 is the positive control template. **B-C)** Example clusters of complex gains across the cell lines shown in Figure 1A, each showing binned purity- and ploidy- transformed read-depth, with the top track showing the associated junction-balanced genome graph (see Methods, [3], with y-axis representing units of per cell copy number (CN) across bins and graph nodes (i.e. intervals). Gray and colored edges represent reference and variant junctions, respectively. Blue edges represent loose ends (see Methods for further details). Bins and junctions are colored as per Figure 1A.

Dewhurst, Yao *et al.* Supplementary Figure 2

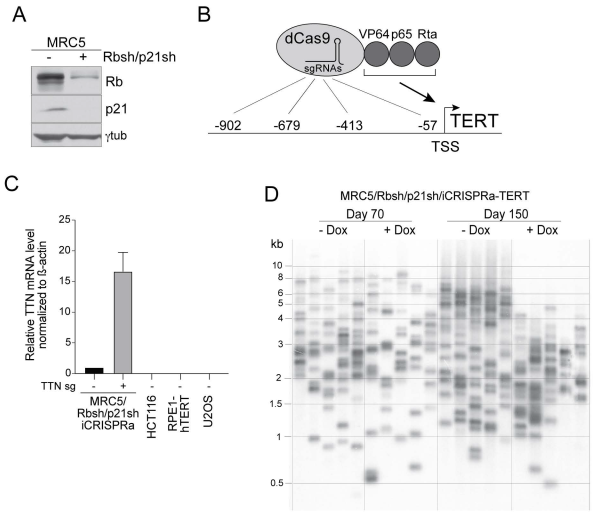


**Supplementary Figure 2. Controlled system for crisis escape. Related to Figure 2.**

**A)** Immunoblot for Rb and 21 in MRC5 cells and MRC5/Rbsh/p21sh cells. **B)** Schematic diagram illustrating the iCRISPRa-TERT system. The positions of the *TERT* activating gRNAs relative to the transcriptional start site (TSS) of the *TERT* gene are shown. **C)** qPCR for *TTN* control gene, activated with a combination of four sgRNAs in MRC5/Rbsh/p21sh iCRISPRa cells (with or without dox, 96 hrs). HCT116, RPE1-hTERT and U2OS cells are included as negative controls. Data are from three independent biological replicates. **D)** STELA of the XpYp telomere in MRC5/Rbsh/p21sh/iCRISPRa-TERT cells after 70 days or 150 days of continuous culture with or without doxycycline. This is a biological replicate of the STELA in Figure 2E.

Dewhurst, Yao *et al.* Supplementary Figure 3

A

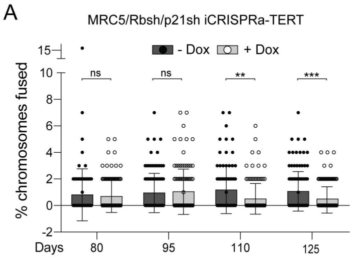


**Supplementary Figure 3. Telomere fusions after controlled escape from crisis. Related to Figure 3.**

**A)** Quantification of percentage of fused chromosomes after the indicated days of continuous culture for MRC5/Rbsh/p21sh/iCRISPRa-TERT cells with and without doxycycline. Data represent means and SDs for three independent biological replicates. *P* values derived from an unpaired Student's t-test. ns, not significant; **, *p* <0.01; ***; *p* <0.001.

Dewhurst, Yao *et al.* Supplementary Figure 4

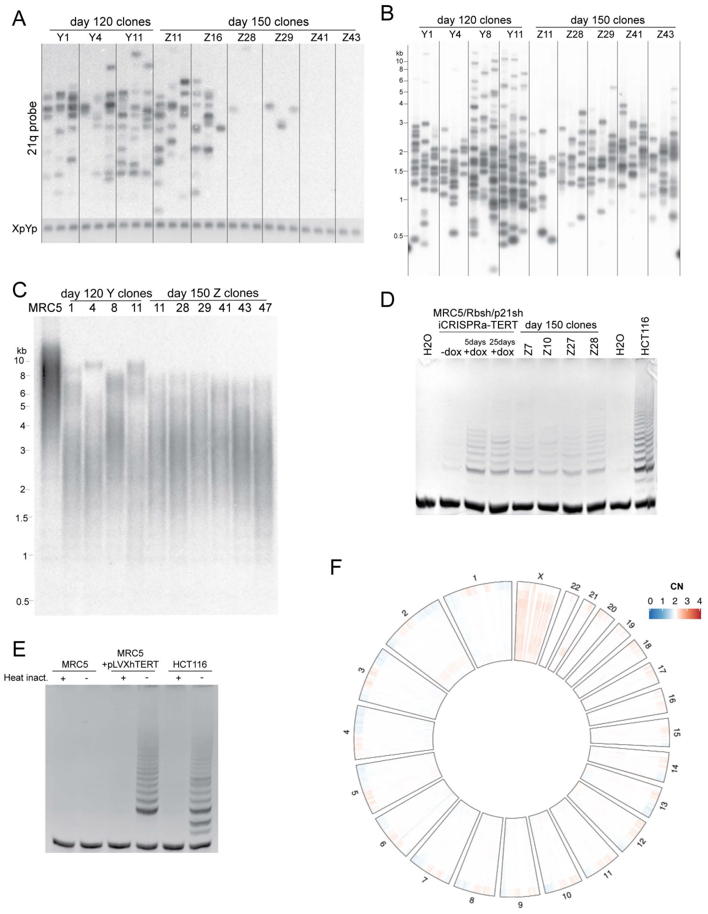


**Supplementary Figure 4. Post-crisis telomere dynamics. Related to Figure 4.**

**A)** Products of telomere fusion PCR on a panel of post-crisis clones (See Figure 4A). Telomere fusions are detected by hybridization to the 21q probe. The control XpYp band is detected with Ethidium bromide staining. **B)** STELA products from a panel of post-crisis clones from both time points. **C)** Telomeric blot on DNA from MRC5 cells and a panel of post-crisis clones from both time points. **D)** TRAP assay showing telomerase activity in MRC5/Rbsh/p21sh/iCRISPRa-TERT treated with doxycycline for the indicated number of days and a selection of post-crisis clones from the day 150 timepoint. HCT116 is included as a positive control. **E)** TRAP assay showing robust telomerase activity in MRC5 cells infected with retroviral pLVX-hTERT. **F)** Circular heatmap showing genome-wide binned purity- and ploidy- transformed read depth (in units of CN across 8 low pass WGS-profiled control CT clones). Heatmap rows correspond to concentric rings in the heatmap.

### Dewhurst, Yao *et al.* Supplementary Figure 5

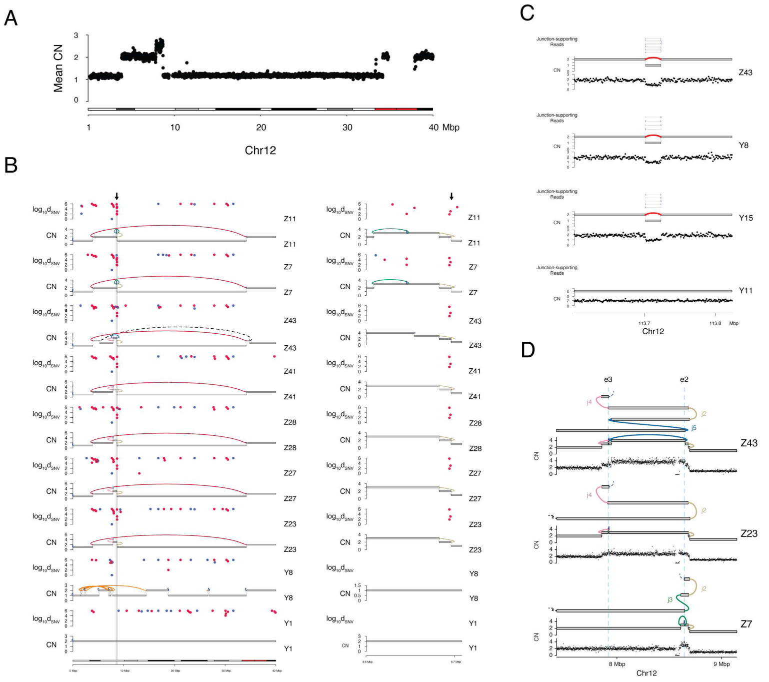


**Supplementary Figure 5. WGS analysis showing genome alterations post crisis clones. Related to Figure 4 and 5.**

**A)** Consensus copy number profile of 47 complex low pass WGS clones targeting chromosome 12p. **B)** Rainfall plot demonstrating SNV patterns as a function of GC vs AT reference nucleotide context, with y-axis showing the logarithm of the inter SNV distance. Highlighted region represents a GC strand coordinated cluster that is found across all four clones in the BFB cluster which harbor junction j2. **C)** Junction supporting read pairs and corresponding drop in coverage at the small deletion on chromosome 12q in representative BFB-like (Z43), chromothripsis-like (Y8), close-relative Y15, and lack of such evidence in distant Y11. **D)** Detailed JaBbA models and linear allele reconstruction at two distinct loose ends, e2 and e3.

Dewhurst, Yao *et al.* Supplementary Figure 6

A

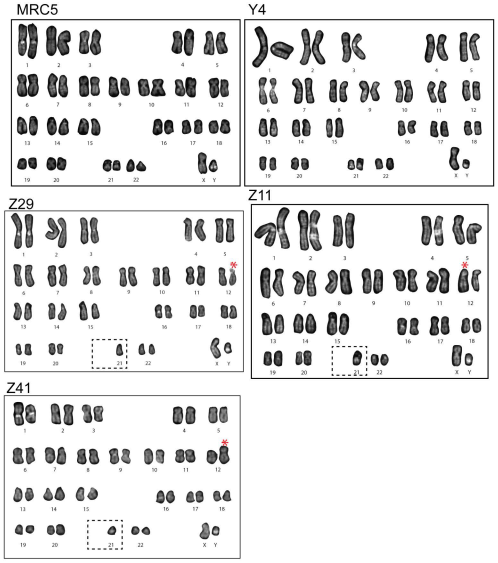


B

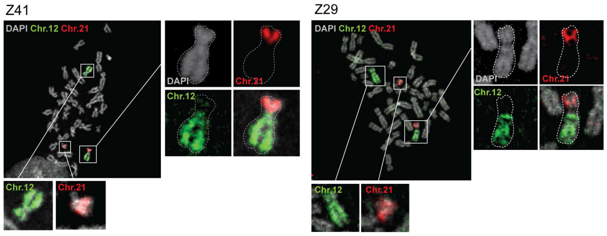


**Supplementary Figure 6. Karyotype evolution in post-crisis clones. Related to Figure 6.**

**A)** DAPI banded karyotypes from the MRC5 parental cell line and a selection of post-crisis clones. Marker chromosomes (chromosome 12) are indicated with a red star. Dashed box indicates the absence of an intact copy of chromosome 21. **B)** Representative images of a metaphase spreads from clones Z41 and Z29 hybridized with both 12 (green) or 21 (red) chromosome paints. DNA stained with DAPI (gray).

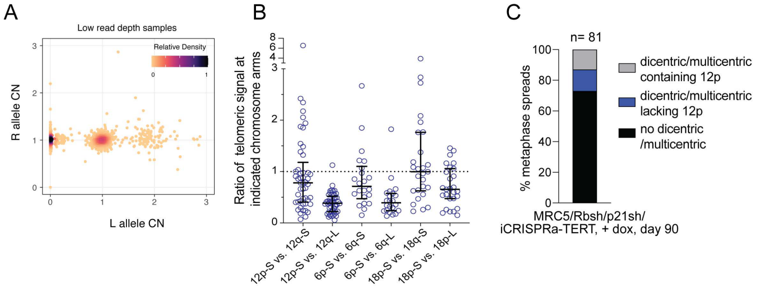


**Supplementary Figure 7. Chromosome 12p allelic imbalance. Related to Figure 7.**

**A)** Scatter plot showing purity- and ploidy-transformed L and R haplotype specific allelic read depth across 12p segments in low pass WGS-profiled post-crisis clones. **B)** Analysis showing that 12p, 6p and 18p arms contain the shortest telomere of those chromosomes. Comparison of the shortest (S) telomere end from each arm (p or q) to each other allele of that chromosome (i.e. q-short allele vs. p-short allele) across chromosome 12, 6 and 18 alleles. Ratio of telomeric intensity from TelG hybridization, as in Figure 7D (S= short allele, L= long allele, based on TelG intensity). Error bars show median and interquartile ranges. Chromosome 12: n= 49 cells, chr.18: n=27, chr.6: n= 22. **C)** Fraction of dicentric or multicentric chromosomes containing 12p in MRC5/Rbsh/p21sh/iCRISPRa-TERT cells in crisis (with doxycycline at day 90) among all dicentric and multicentric chromosomes. Based on analysis of images as in Figure 7E. n=81 metaphases scored from two independent experiments.

**Supplementary Table 2: Number of clones analyzed by high and low-pass WGS**

Summary of the number of clones analyzed by WGS (high and/or low pass) from each time point.

**Supplementary Table 3: Oligos used in this study**

Sequences of all oligos used in this study (see also Methods).

**Supplementary Table 1: SV40T immortalized post-crisis cell lines**

| Name | Cell Line/Tissue Source | Transformation | Telomerase status | Reference | Kind gift of |
|---|---|---|---|---|---|
| HA-1M PD216 | Human Embryonic Kidney cells | SV40 | Positive (see ref.) | [29] | Silvia Bacchetti/AdVec |
| SW13 PD184 | IMR90 lung fibroblasts | SV40 | Positive (Supp. Fig.1) | [28] | Jerry Shay, UTSW |
| SW26 PD130+ | IMR90 lung fibroblasts | SV40 | Positive (Supp. Fig.1) | [28] | Jerry Shay, UTSW |
| Bet3B p25 post-crisis | NHBE-10 bronchial epithelial cells | SV40 | Positive (see ref.) | [27] | Roger Reddel, CMRI |
| Bet3K p25 post-crisis | NHBE-10 bronchial epithelial cells | SV40 | Positive (see ref.) | [27] | Roger Reddel, CMRI |
| BFT3B p28 post-crisis | BF-10 bronchial fibroblasts | SV40 | Positive (see ref.) | [27] | Roger Reddel, CMRI |
| BFT3K p34 post-crisis | BF-10 bronchial fibroblasts | SV40 | Positive (see ref.) | [27] | Roger Reddel, CMRI |

**Supplementary Table 2: Number of clones analyzed by high and low-pass WGS**

| Name | Number Sequenced | |
|---|---|---|
| | Low Pass | High Pass |
| Parental cell lines | 3 | 1 |
| Day 120 (Y) clones | 37 | 5 |
| Day 150 (Z) clones | 83 | 8 |
| Control (CT) clones | 8 | 0 |

**Supplementary Table 3: Oligos used in this study**

| Oligo name and sequence | Source |
|---|---|
| TERT_gRNA_1- AGTCGCGGGGAAGTGTTGCA | This study |

| | |
|---|---|
| TERT_gRNA_2- ATCTGCCAGACAGAGTGCCG | This study |
| TERT_gRNA_3- TCGAATCGGCCTAGGCTGTG | This study |
| TERT_gRNA_4- GAAACTCGCGCCGCGAGGAG | This study |
| TTN_gRNA_1- CCTTGGTGAAGTCTCCTTTG | [32] |
| TTN_gRNA_2- ATGTTAAAATCCGAAAATGC | [32] |
| TTN_gRNA_3- GGGCACAGTCCTCAGGTTTG | [32] |
| TTN_gRNA_4- ATGAGCTCTCTTCAACGTTA | [32] |
| TERT qPCR forward- GGAGCAAGTTGCAAAGCATTG | This study |
| TERT qPCR reverse- TCCCACGACGTAGTCCATGTT | This study |
| TTN qPCR forward- TGTTGCCACTGGTGCTAAAG | This study |
| TTN qPCR reverse- ACAGCAGTCTTCTCCGCTTC | This study |
| β-actin qPCR forward-TGGATCAGCAAGCAGGAGTATG | This study |
| β-actin qPCR reverse- GCATTTGCGGTGGACGAT | This study |
| STELA telorette1- TGCTCCGTGCATCTGGCATCCCCTAAC | [34] |
| STELA telorette2- TGCTCCGTGCATCTGGCATCTAACCCT | [34] |
| STELA telorette3- TGCTCCGTGCATCTGGCATCCCTAACC | [34] |
| STELA telorette4-TGCTCCGTGCATCTGGCATCCTAACCC | [34] |
| STELA telorette5- TGCTCCGTGCATCTGGCATCAACCCTA | [34] |
| STELA telorette6- TGCTCCGTGCATCTGGCATCACCCTAA | [34] |
| STELA XpYpE2- TTGTCTCAGGGTCCTAGTG | [34] |
| STELA teltail - TGCTCCGTGCATCTGGCATC | [34] |
| STELA XpYpB2- TCTGAAAGTGGACCTATCAG | [34] |
| Fusion PCR XpYpM- ACCAGGTTTTCCAGTGTGTT | [35] |
| Fusion PCR 17p6- GGCTGAACTATAGCCTCTGC | [35] |
| Fusion PCR 21q4- GGGACATATTTTGGGGTTGC | [35] |
| Fusion PCR XpYpc2tr- GCTATGGCTTCTTGGGGC | [35] |
| Fusion PCR 21q-seq-rev2-ACACAGAAGGTTGATATACACAG | [35] |
